# Combinatorial mutagenesis of N-terminal sequences reveals unexpected and expanded stability determinants of the *Escherichia coli* N-degron pathway

**DOI:** 10.1101/2025.05.22.655665

**Authors:** Sabyasachi Sen, Nastassja Corrado, Alexander R. Tiso, Khai Khee Kho, Aditya M. Kunjapur

## Abstract

Although it is known that residues near the N-terminus can influence protein stability, there has not been a comprehensive effort to document how these penultimate residues impact protein stability in prokaryotes. Here, we utilize combinatorial mutagenesis, cell sorting, and next generation sequencing to perform the deepest sequence coverage screen of the bacterial N-degron pathway of proteolysis. We present nuance and exceptions to the N-terminus (P1) functioning as a primary stability determinant. We reveal stability contributions for P2-P5 motifs, including lowered stability for clustered bulky residues and Gln, and heightened stability for negatively charged residues, Pro, and Gly. We find that P1 Cys can be an N-degron component in a sequence-specific manner. Furthermore, we employ stability-predictive machine learning to identify motifs with unexpected fates. Our work expands the stability determinants of the N-degron pathway with unprecedented granularity, serving as a resource for N-degron identification, N-degron design, and future molecular basis elucidation.

## Introduction

Protein degradation plays a key role in supporting natural and engineered functions of bacteria. Fundamentally, protein degradation enables the recycling of proteins that have either transient, stimuli-responsive lifespans or are aberrantly expressed^1^. This phenomenon intersects with human health in myriad ways. For example, certain bacterial proteins that are directly involved in protein degradation, such as the Clp protease system in the pathogen *Mycobacterium tuberculosis*^2–4^, are attractive antibiotic targets. Additionally, inducible degradation of essential proteins using BacPROTACs^5^ has been performed to guide antibiotic target discovery. In applied biotechnology efforts, protein degradation tags are frequently used to attenuate hysteresis in genetic circuits within industrial workhorse strains such as *Escherichia coli*^6–8^.

One impactful protein degradation system is the N-degron pathway, which modulates protein half-life across minutes to hours. The pathway relies on sequence-specific N-terminal binding interactions to initiate degradation and is conserved across both prokaryotes and eukaryotes^9^. In *E. coli,* bulky (Leu, Phe, Trp, Tyr) or positively charged (Arg, Lys) amino acids at the N-terminus have been identified as a primary signal for N-degradation^10,11^. The N-recognin ClpS, a prokaryotic homolog of the eukaryotic N-recognin Ubr1^12^, recognizes and binds to N-degron-tagged proteins that have bulky amino acids at the N-terminus and deposits them at the ClpAP protease to substrate turnover^13–16^. A second key *E. coli* protein, leucyl-phenylalanyl transferase (LFTR), recognizes positively charged amino acids at the N-terminus and will append one or more Leu/Phe to that residue to generate a new ClpS substrate^17–19^.

Despite these reported N-terminal patterns, recent work has shown that the identity of the N-terminal (P1) residue is not a definitive determinant of protein fate. Residues through at least the first 5 positions of a protein (P1-P5) have been shown to impact stability through the pathway, including situations where the fate of a protein appears to defy its N-terminal residue^16,18,20–25^. This is consistent with observations that downstream residues influence the dissociation constants of purified N-terminal binding proteins, including ClpS, when they are repurposed for next-generation protein sequencing applications^26–29^. These important yet limited observations strongly motivate a comprehensive analysis of N-degron sequence stability determinants in bacteria (Figure 1A). The benefits of such an analysis would impact various disciplines. For example, synthetic and chemical biologists have increasingly designed alternative protein N-termini for non-degradative purposes such as bio-orthogonal conjugation reactions^30–33^, live cell cleavage of N-terminal tags or signal peptides, or protein ligation or excision via inteins, sortases, or subtiligases^34–38^. Additionally, in the fields of *de novo* design and protein engineering there is a need for comprehensive experimental datasets that profile how synthetic sequences impact *in vivo* protein stability^39^. For these applications and more, it is valuable to understand when a generated neo-N-terminus can form an N-degron.

**Figure 1.**
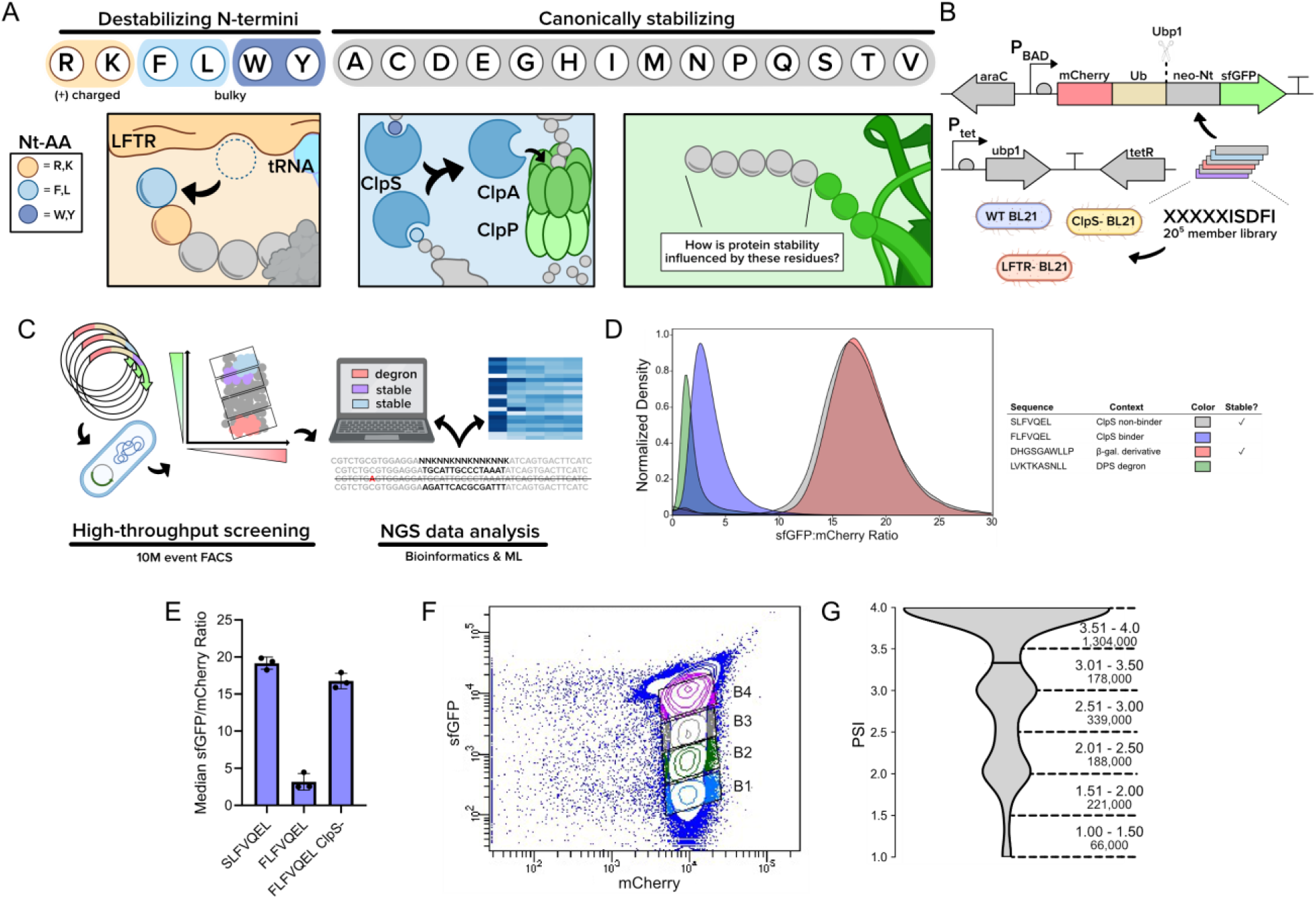
Overview and HTS workflow for N-terminal sequence analysis. **A.** An overview of the *Escherichia coli* Leu/N-degron pathway. The top row shows the expected fate for proteins with the listed amino acids at the N-terminus, where the orange and blue shaded residues are canonically destabilizing. The left-most box illustrates the leucyl/phenylalanyl transferase LFTR, appending Phe or Leu onto an N-terminal Arg or Lys. The middle box illustrates the ClpS adaptor protein that binds N-terminal F, L, W, and Y and recruits the protein to the ClpAP protease for degradation. The right-most box shows the dual reporter after the action of a ubiquitin (Ub) cleavase, revealing a neo-N-terminus (neo-Nt) on an sfGFP reporter. **B.** Genetic constructs for the protein stability assay. The first plasmid harbors a synthetic gene fusion that encodes the dual fluorescent reporter under control of an arabinose-inducible promoter. The second plasmid harbors a gene encoding ubiquitin cleavase from *Saccharomyces cerevisiae* under control of a tetracycline-inducible promoter. Constructs were tested in wild-type BL21 as well as BL21 derivatives deficient in *clpS* and *aat* (LFTR). **C.** The HTS protein stability workflow used in this study. **D.** sfGFP:mCherry ratio density histograms for over 100,000 events collected for cells expressing Ubp1 as well as literature control N-terminal sequences embedded in a dual fluorescent reporter construct. Stable sequences were SLFVQEL and DHGSGAWLLP and unstable sequences were FLFVQEL and LVKTKASNLL. **E.** Known ClpS-binding degron FLFVQEL returns stabilizing sfGFP:mCherry ratio values in the absence of ClpS. Data shown is median fluorescence intensity ± standard deviation for at least 100,000 events captured across clonal biological triplicates. **F.** FACS binning of the XXXXXISDFI library within wild-type BL21 **G.** Violin plot showing the PSI distribution of 2.30M collected sequences from the wild-type BL21 dataset. The number of sequences is rounded to the nearest thousand. Figures 1A-1C generated using Biorender content.

To date, the characterization of the bacterial N-degron pathway has been performed using low-throughput protein half-life assays, *in vitro* pathway reconstitution, and affinity purification to screen dozens to hundreds of putative N-degrons at once^10,17,18,22,24,40,41^. While these methods have pioneered our understanding of N-degrons, they have their limitations. Pulldown assays are limited to sequences enriched within the bacterial proteome. Low-throughput reporter screens require extrapolation from smaller datasets. Finally, *in vitro* experimentation can sometimes present artifacts or deviations from *in vivo* pathway behavior. To address these limitations, recent advances within the protein degradation field, including exemplary work by Timms et al.^42^, have leveraged high-throughput screening and analysis of N-terminal libraries derived from the human proteome to analyze roughly 10^4^ sequences within human cells. Similar approaches have been taken to study protein degradation^43–46^, however no reported studies to date have screened large DNA libraries to characterize prokaryotic branches of the N-degron pathway.

To investigate this knowledge gap, we evaluated the role of the five N-terminal amino acids using combinatorial mutagenesis, a reporter protein assay in wild-type and N-recognin-deficient bacterial hosts, fluorescence-activated cell sorting (FACS), next-generation sequencing (NGS), and bioinformatic analysis. From over 2.2 million unique sequences screened and collected, we have mapped the N-degron pathway with increased granularity. We share heatmaps that reveal P2-P5 preferences for canonically destabilizing P1 residues. We additionally reveal a series of sequence-based stability determinants. Specifically, at P2, Gln (destabilizing), Pro (stabilizing), and Gly (stabilizing) show the largest single-residue stability shifts, exceeding select destabilizing P1 residues. In an analysis of low stability sequences without the canonically destabilizing P1 FLYWRK, we found motifs rich in P1 Gln, Cys, or His with bulky and negatively charged residues in P2-P5. Furthermore, additive stability effects were observed for motifs rich in bulky residues (destabilizing), negatively charged residues (stabilizing), and flexible Gly/Ser residues (stabilizing). We then trained a machine learning model, N-FIVE, on our dataset and harnessed it to predict sequences that break canonical stability rules. In total, our manuscript serves as an in-depth resource for understanding N-degron pathway specificity and designing bacterial degrons.

## Results

### Design, validation, and library sorting using an *in vivo* N-degron pathway fluorescent reporter

To monitor *in vivo* protein degradation, we adapted the ubiquitin reference^47^ and global protein stability techniques^48^ by creating an mCherry-ubiquitin-degron-sfGFP-His6x genetic fusion that was cloned into a plasmid (p15a ori, CmR, P_araBAD_). Ubiquitin (Ub), considered inert within *E. coli*, is scarlessly cleaved at its C-terminus upon the expression of Ub protease Ubp1 that is housed on a second plasmid (ColE1 ori, KanR, P_tet_). This cleavage simultaneously reveals a neo-N-terminus (neo-Nt) that is attached to sfGFP while producing a stable fluorescent mCherry-ubiquitin motif. The subsequent degradation of sfGFP that are fused to an N-degron should result in lower sfGFP:mCherry ratios for pathway substrates relative to stable neo-Nt-sfGFP proteins. This dual fluorescent reporter assay avoids various confounding effects, including variable protein expression due to the 5’ ORF mRNA sequence or the amino acid sequence near the translational start site^49–57^. We selected *E. coli* BL21 as the expression host due to its deficiency of the Lon and OmpT proteases, helping to minimize proteolytic crosstalk and to isolate the effect of the ClpSAP system. To further isolate the impact of each key degradation adaptor protein, we generated two separate variants of BL21 where either ClpS or LFTR were inactivated through the insertion of in-frame stop codons using multiplex automatable genome engineering^58,59^. These genetic constructs and assay design serve as the experimental basis for our high-throughput screening platform (Figure 1B, Figure 1C).

To verify assay functionality, we screened neo-Nt sequences previously reported to be either strong or non-interactive ClpS/N-degron pathway substrates. For positive controls, we selected reported ClpS ligand FLFVQEL^24,60^ and LVKTKASNLL, the latter derived from ClpSAP substrate Dps^18^. For negative controls, we selected SLFVQEL, a known ClpS non-interacting sequence, and DHGSGAWLLP^10^, a reported stable motif derived from the first 10 amino acids of β-galactosidase. Using flow cytometry, we observed unimodal distributions of sfGFP/mCherry ratio across all tested degrons and we selected median sfGFP/mCherry ratio as a comparative metric. Ratiometric comparisons revealed an average 6.0 and 5.9-fold dynamic range between SLFVQEL/FLFVQEL and DHGSGAWLLP/LVKTKASNLL sets, respectively (Figure 1D). We additionally screened FLFVQEL in the *clpS^−^* host and observed a 5.2-fold ratio increase over the wild-type strain, further evidencing that observed ratiometric changes from our assay matched expectations (Figure 1E). When we screened our control sequences on a BD FACS Aria II, we observed order of magnitude separation between N-degrons and stable neo-Nt fusions (Figure S1A). Towards deep sequence profiling, we first validated our N-degron screening platform using a 60-member library, where we profiled the P1-P3 stability of eukaryotic-derived N-degrons containing Nt-Arg-Cys^61^. From this experimentation, we identified a strongly stabilizing influence of acidic residues at P3 that we verified *via* Western blot.

We next sought to expand our library size to determine the stability contributions for all combinations of the first five amino acids through the N-degron pathway. We selected a sequence template derived from the eukaryotic protein RAP2.2 (CGGAIISDFI)^62^ due to its consistent behavior and high dynamic range between the aforementioned sequence (stable) and a variant that contains an N-terminally appended Arg (RCGGAIISDFI, unstable). Using five consecutive NNK codons located at the neo-N-terminus of the reporter (XXXXXISDFI, theoretical library size: 3.2E6), we transformed the library into unmodified, wild-type (WT) and knockout BL21 strains carrying a Ubp1-expressing plasmid. We then induced expression of both the reporter and Ubp1 and subsequently collected 10M gated events using fluorescence activated cell sorting (FACS). We consistently observed most cells within the population in a large, high-sfGFP group with a tail that extended across orders of sfGFP fluorescence magnitude (Figure S1B). We sorted mCherry-expressing cells into four bins that covered the majority of the sfGFP fluorescence range (Figure 1F). Upon analyzing 100-150M reads of an amplicon containing the mutagenized region from the WT, LFTR^−^, and ClpS^−^ sorts, we obtained 2.29M, 2.34M, and 2.19M unique sequences per host, respectively. To evaluate the stability of each collected sequence, we then calculated the Protein Stability Index (PSI), a weighted average utilized in high-throughput protein degradation experimentation^42,48,46,63^. Here, 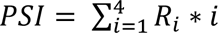, where *R_i_* is the fraction of reads present in bin *i.* In line with expectations that most P1-P5 sequences are stable, PSI distributions were skewed towards high stability values, with over 82% of wild-type sequences belonging in the top half of the PSI range (Figure 1G).

### Deep sequence profiling reveals N-degron specificity through the first five amino acids

To study the substrate-sequence preferences and contributions of various N-degron pathway components, we generated average PSI heatmaps for all amino acid and position combinations for wild-type (Figure S2A), LFTR^−^ (Figure S2B), and ClpS^−^ hosts (Figure S2C). Here, the expectation is that amino acid motifs with lower PSI are recognized *in vivo* by LFTR and/or ClpS, which initiate sfGFP degradation. To visualize how individual amino acids downstream of P1 impact stability while minimizing expression-based variability, we computed the average PSI for each AA-position combination in the P1-P5 space and plotted the differences between WT and ClpS^−^ datasets (Figure 2A). The most distinct shifts in PSI were based on the P1 residue, in line with the expected role of P1 as a critical stability determinant. The order for destabilizing P1 residues is roughly tiered from lowest to highest average PSI as Phe = Arg < Leu < Trp = Lys = Tyr. An analysis of the 100,000 lowest PSI sequences revealed an enrichment for five of these six residues (excluding Lys) at and near P1 (Figure S2D). In a comparison of P1 residues between the wild-type and LFTR^−^ datasets, there was a linear correlation of average PSI values with an r-squared of 1.0 for the 17 P1 amino acids that are expected to behave similarly (Figure S2E). As expected, Arg and Lys at P1 exhibited differential stability between LFTR^−^ and WT hosts with lower average values in the WT dataset. In this analysis, P1 Pro was excluded due to confounding effects when it is not uniformly cleaved by Ulp1; P1 Pro is the only known instance where fused ubiquitin is not fully excised by Ubp1^64^. In line with the reported role of ClpS as a sequence-specific binder and degradation initiator, the ClpS^−^ dataset shows sequence-agnostic parity in the absence of the N-recognin, with a small bias for Arg, Leu, Pro, and Val at low PSI (Figure S2F). All cells in the corresponding ClpS^−^ P1-P5 heatmap are within 0.17 PSI units of each other, demonstrably smaller than the >1 PSI differences observed between canonically stable and unstable P1 residues in WT BL21 and LFTR^−^ BL21 (Figure S2A and S2B).

**Figure 2.**
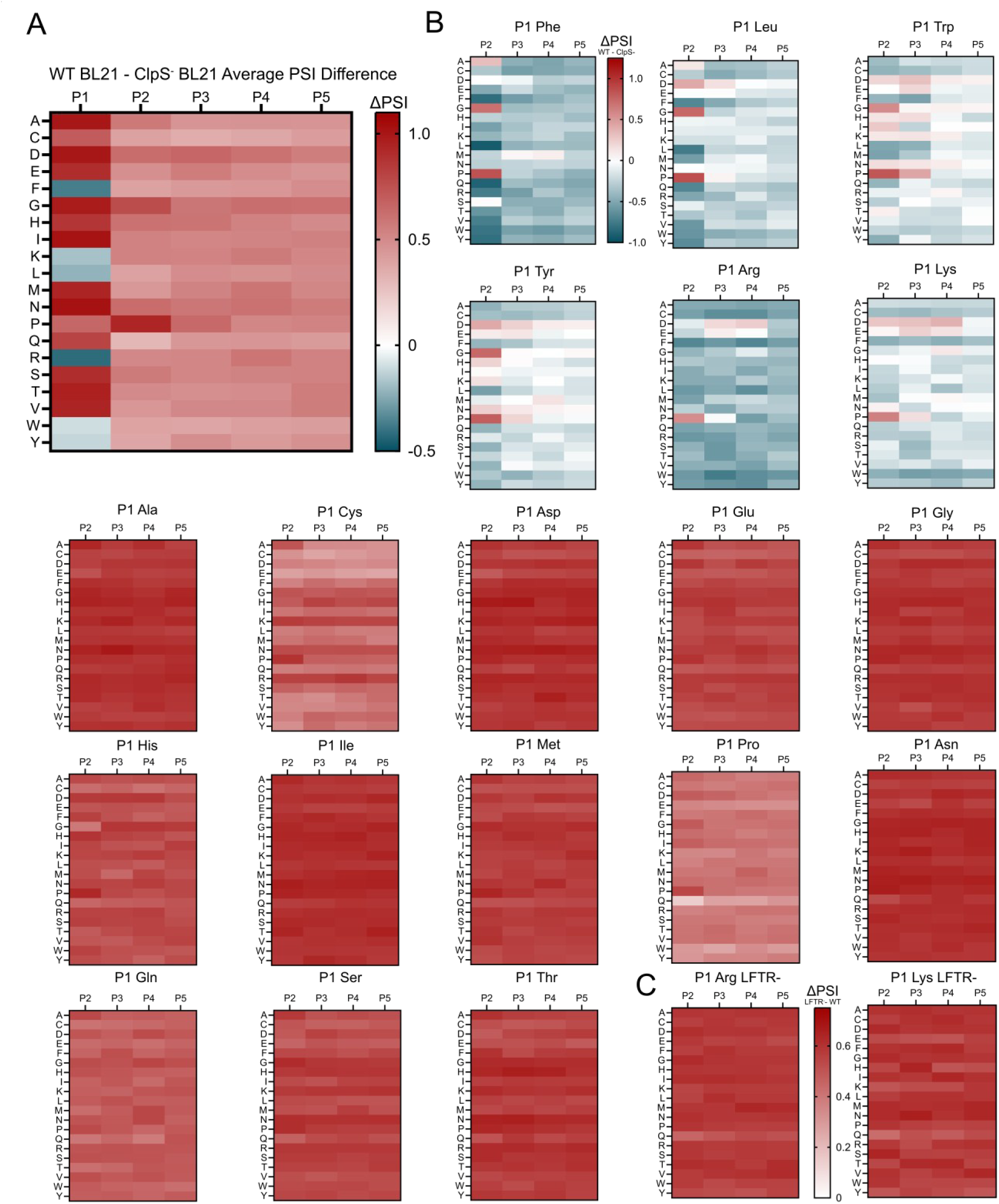
Summary of stability contributions for penultimate amino acids. **A.** Heatmap visualizing the bulk P1-P5 preferences of the *Escherichia coli* N-degron pathway visualized as the average change in PSI between wild-type BL21 and ClpS^−^ BL21 datasets for P1-P5 amino acid-position combinations. **B.** Heatmaps visualizing P2-P5 preferences each standard amino acid at P1 using the average change in PSI between wild-type BL21 and ClpS-BL21 for P2-P5 amino acid-position combinations. **C.** Increased P2-P5 stability profile for P1 Arg and Lys in *Escherichia coli* deficient in LFTR visualized using the average change in PSI between LFTR^−^ BL21 and WT BL21 datasets for P2-P5 amino acid-position combinations.

Next, we visualized the P2-P5 preferences for all P1 residues by calculating the average PSI difference between WT and ClpS^−^ datasets for all amino acid-position combinations (Figure 2B). We observed PSI variability at P2-P5 for sequences with a canonically destabilizing P1. Specifically, while most 2-position sequence combinations led to PSI decreases, there were instances where the drops in PSI varied in magnitude. For example, destabilizing PSI shifts with P1 Arg were smaller when negatively charged residues were present particularly in P3 & P4, as well as Pro in P2. In parallel, P1 Phe showed smaller shifts for small residues (Ala, Gly, Ser) in P2-P5 and Pro in P2. Frequently, the presence of a bulky residue in P2 lead to greater PSI drop magnitudes. P1 Leu, Trp, Lys and Tyr all exhibit PSI differences between different amino acid combinations, indicating unique P2-P5 preferences for these canonically destable P1 residues. These results further evidence that P1 identity cannot be viewed in as an exclusive stability determinant, particularly when identifying highly destabilizing N-degrons.

We next profiled the specificity of LFTR by analyzing the PSI difference between WT and LFTR-datasets. In a comparison of average PSIs values between the WT and LFTR- dataset, a clear PSI difference was observed for P1 Arg and Lys values, suggesting increased stability in the absence of LFTR (Figure S2G). When analyzing the shift in PSI at P2-P5 for P1 Arg and Lys, a clear and uniform increase in PSI is observed (Figure 2C). Arg and Lys residues at P2-P5 did not appear to influence stability, suggesting that internal residues were not contributing to destabilization in bulk (Figure S2H & S2I). One exception was for FRXXX sequences, where P2 Arg led to the largest decrease in mean PSI within the subset of sequences with internal Arg residues. In total, Arg and Lys do not notably impact stability in the absence of LFTR, as expected.

### P2 residues including Pro, Gly, Gln, and Ala are critical stability determinants

Having shown that residues adjacent to P1 can have a pivotal role in stability determination, we next focused on key stability determinants at P2. As P2 is positioned on the periphery of the ClpS binding pocket in crystallized bound peptide ligands, docking studies have not clearly elucidated a role for this position (Figure S3A). Within our datasets, several amino acids showed stabilizing effects at this position. We observed this most distinctly from Gly and Pro, with the effect’s magnitude having a maximal impact at P2 and waning with distance from P1. Commonly enriched in stable, PSI >3 motifs (Figure S3B), these two residues had the highest PSI change in the WT - ClpS^−^ heatmap. P2 Pro paired with P1 FLWYRK led to a significant 1.05-unit increase in PSI (p <1E-99, ES = 0.71) (Figure 3A). Notably, P2 Pro often overrode the stability contribution of canonically destabilizing P1 residues, with larger groups being observed in high stability ranges (Figure 3B). Only select P1 Arg or P1 Lys motifs were notably present at lowered PSI values relative to the rest of the P1 dataset. A similar, but smaller in magnitude 0.63-unit PSI increase was observed for P2 Gly (p<1E-99, ES = 0.47) (Figure 3C). Upon testing clonal isolates with P2 Gly, we found motifs that stabilize P1 Leu and Trp. Interestingly, P1 Arg remained destabilizing, potentially due to Gly having an extended distance from the neo-P1 upon LFTR activity (Figure 3D).

**Figure 3.**
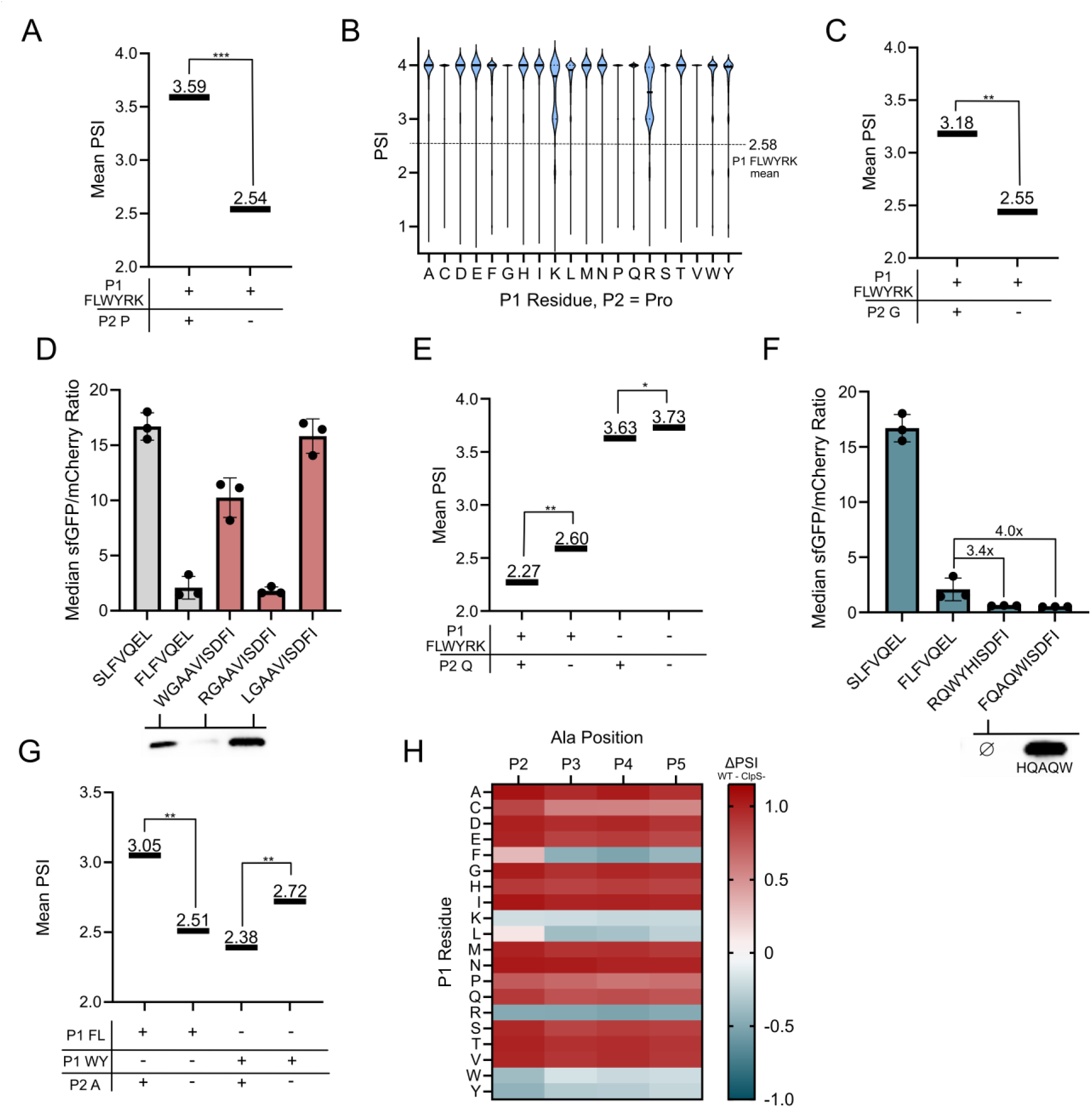
P2 residues including Pro, Gly, Gln, and Ala are critical stability determinants. **A.** Stability increase from P2 Pro being paired with a canonically unstable N-terminus. Mean PSIs for wild-type sequence subsets where the number of sequences per condition are (++): 33,674, (+−): 750, 888; (++/+−) ES: 0.71. **B.** Violin plot showing PSI distributions for various P1 residues with P2 Pro in the WT dataset. The mean PSI for all P1 FLWYRK residues is shown as a dashed line. **C.** Stability increase from P2 Gly being paired with a canonically unstable N-terminus. Data shown is average PSI for wild-type sequence subsets where the number of sequences per condition are (++): 41,975, (+−): 742,587; (++/+−) ES: 0.47. **D.** Clonal P2 Gly isolates show stabilizing effects for two non-LFTR modified N-terminal motifs. Flow cytometry data for various P2 glycine motifs (top). Western blots of hardcoded sequences (bottom). **E.** An amplified stability decrease when P2 Gln is paired with a canonically unstable N-terminus. Data shown is average PSI for wild-type sequence subsets where the number of sequences per condition are (++): 29,982, (+−): 754, 580, (−+): 52,021, (−−): 1,459,940; (++/+−) ES: 0.26. **F.** Clonal P2 Gln isolates show decreased stability measurements relative to a literature control. Flow cytometry data for various P2 glutamine motifs (top). Western blots of hardcoded sequences (bottom). No band was visible for the FQAQWISDFI sequence, indicated by ∅. **G**. P2 Ala stabilizes P1 F or L and further destabilizes P1 W or Y. Data shown is average PSI for wild-type sequence subsets where the number of sequences per condition are (+−+): 14,127, (+−−): 272,399, (−++): 12793, (−+−): 232,067; (+−+/+−−) ES: 0.44, (−++/−+−) ES: 0.27. **H**. Bulky P1 stability shifts from Ala are primarily when Ala is at P2. Heatmap depicting the change in PSI between WT and ClpS^−^ datasets for all P1 residues with Ala at a given P2-P5 position. p-values were evaluated using a Mann-Whitney U test. Effect size magnitude (ES) ranges from 0 to 1, as evaluated using Cliff’s Delta. * = p <0.05, ES 0-0.25; ** = p <0.05, ES 0.25-0.50, *** = p <0.05, ES 0.50-0.75, **** = p <0.05, ES 0.75-1.0. Comparisons are abbreviated as (P1 FLYWRK presence, P2 presence) or (P1 FL presence, P1 WY presence, P2 presence).

Gln at position 2 (and to a lesser extent at P3-5) is amongst the most destabilizing P2 amino acids in both WT and LFTR^−^ BL21, providing a −0.51 PSI unit decrease at P2 in the WT dataset. Fascinatingly, Gln at P2 is just as strong a determinant of protein degradation as L at P1 (−0.52 units). The destabilizing effect was amplified when Gln was paired with P1 LFWYRK (p <1E-99, ES = 0.26), showing a 3.2x decrease in the difference in mean PSI compared to Gln paired with an alternative P1 residue (Figure 3E). The stability impact of Gln was visible across WT and LFTR^−^ datasets and absent in the ClpS^−^ dataset, suggesting a ClpS-mediated preference (Figure S2A and S2C). We sought to validate this observation using clonal examples. Upon pairing Gln with a destabilizing P1 residue, we identified sequences that resulted in sfGFP:mCherry ratios that were 3.4 and 4.0 times lower than our degron control (RQWYHISDFI : FLFVQEL and FQAQWISDFI : FLFVQEL, respectively). Furthermore, neither FQAQWISDFI nor RQWYHISDFI showed no visible degron-sfGFP band when visualized via Western blot (Figure 3F, Supplementary File 2), suggesting strong turnover of the construct.

A final residue of note was P2 alanine. Specifically for bulky destabilizing P1 residues, P2 Ala can have either a stabilizing or destabilizing effect depending on the P1 residue. For P1 Phe or Leu, P2 Ala leads to a significant increase in PSI compared to other P2 residues (p<1E-99, ES = 0.43). Furthermore, for P1 Trp or Tyr, P2 Ala provides a significant destabilizing effect (p<1E-99, ES = 0.27). This observation reveals a differential specificity amongst bulky destabilizing P1 residues (Figure 3G). This effect was only observed for Ala at P2 and not P3-P5 for those specific P1 residues (Figure 3H).

We additionally examined whether the native *E. coli* methionine aminopeptidase (MetAP) could generate an N-degron by excising Met at P1 to generate a degradation-promoting motif starting at P2. We compared the PSI for motifs carrying Nt-Met (MXYYY) to those same motifs when they appear in the first four positions (XYYYZ) within the wild-type dataset. We observed that residues that are canonically destabilizing at the X-position had increased mean PSI values in MXYYY motifs when compared to XYYYZ motifs (Figure S3C & Figure S3D). In line with the published specificity of MetAP^65,66^, it did not appear that Met-L/F/W/Y/R/K motifs produced low PSI N-degrons in bulk.

### Destabilizing motifs include unexpected P1 residues Cys, His, and Gln with specific P2-P5 patterns

Having found that P2 residues can drastically influence stability, we next investigated the stability contributions of all 400 P1/P2 pairs in greater detail. First, we observed that LFWYRK at P1/P2 were highly represented in low stability sequences. In an analysis of the 100 most enriched P1-P2 motifs at low PSI, 90/100 contained at least one FLWYRK residue, and 34/36 possible combinations containing two FLWYRK residues were present, showing an overrepresentation of these six residues within destabilizing motifs relative to other combinations at P1/P2 (Figure 4A). Enriched residues not in FLWYRK within the low stability P1/P2 space of select residues including P1 Cys, P1 and P2 Gln, and P2 Trp (Figure 4B). Furthermore, when visualizing all P1/P2 combinations that were enriched at low PSI but did not carry a canonically destabilizing P1 residue, Gln was overwhelmingly present (Figure S4A). From this finding, we next investigated the sequence profiles for low stability sequences with canonically stable P1 residues.

**Figure 4.**
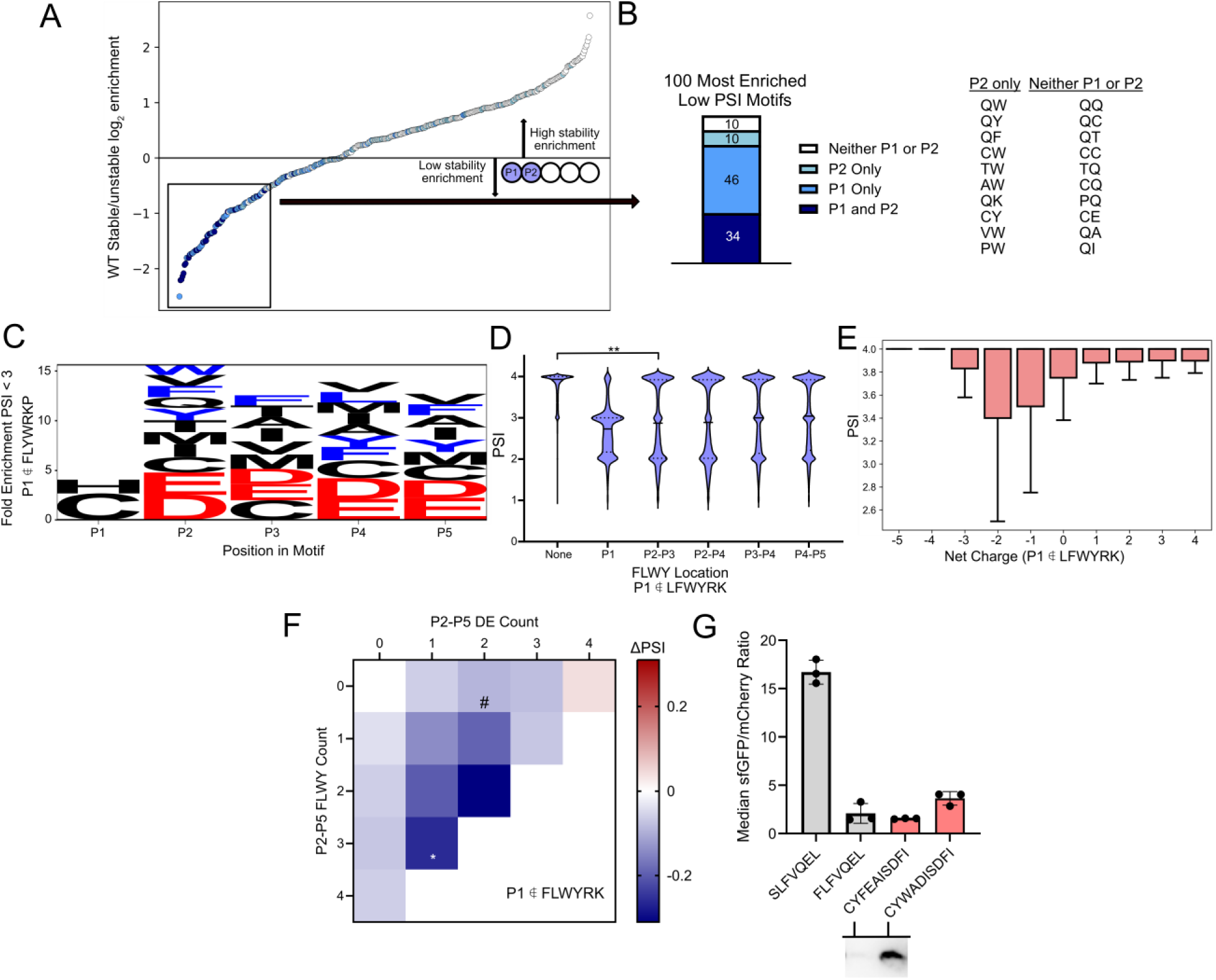
Destabilizing motifs can contain P1 Cys and Gln with specific P2-P5 motifs. **A.** Log_2_ enrichment ratio plot of the 400 possible stable (PSI >3)/unstable (PSI<2) P1-P2 dipeptide motifs. Sequences containing LFWYRK at various positions are listed in the following pairs: (P1 and P2, dark blue), (P1 only, light blue), (P2 only, light green), (neither P1 or P2, white). **B.** LFWYRK composition of the 100 lowest PSI P1/P2 motifs. Motifs with P1 not in FLWYRK are listed below the plot. **C.** Low stability sequences with canonically stable N-termini are rich in negatively charged and bulky amino acids in P2-P5. Weblogo depicting fold enrichment amino acids in low PSI (PSI <3) motifs that do not contain FLWYRKP at P1. Bulky residues (FLWY) depicted in blue and negatively charged residues (DE) depicted in red. **D.** Multiple bulky residues lower mean PSI. Violin plot of wild-type BL21 data showing PSI distributions for sequences with bulky residues at various positions. A comparison between motifs with no bulky residues (n = 1,511,962) and motifs with bulky residues only in P2 and P3 (n=56,341) shows a significant difference between mean PSI values (p < 0.05, ES = 0.25) **E.** PSI boxplot for sequences with varying net charge and canonically stable P1 residues shows the opposite charge trend to low stability P1 FLWYRK sequences. P1 Pro is excluded. **F.** For WT sequences with canonically stable P1, the largest PSI drops are observed for motifs with both bulky and negatively charged P2-P5 residues. Change in PSI is relative to the 0 P2-P5 FLWY, 0 P2-P5 DE dataset (mean PSI = 3.78). All p-values relative to (0,0) are p <0.05 except cells marked with # which are p >0.05. * = ES > 0.25. **G.** Identification of an Nt-Cys degron. (Top) flow cytometry data, (bottom) Western blots of the degron-sfGFP product for candidate degrons with N-terminal cysteine. p-values evaluated using a Mann-Whitney U test, effect size magnitude (ES) ranges from 0 to 1, as evaluated using Cliff’s delta. * = p <0.05, ES 0-0.25; ** = p <0.05, ES 0.25-0.50, *** = p <0.05, ES 0.50-0.75, **** = p <0.05, ES 0.75-1.0

We then analyzed moderate to low stability (PSI < 3) sequences without LFWYRK at P1. In this analysis, Cys and His were enriched at P1. Additionally, bulky residues (LFWY), acidic residues (DE), and assorted other residues (MVC) were enriched at P2-P5 (Figure 4C). Next, a series of subsets were analyzed. First, as previous studies have reported that bulky residues downstream of the N-terminus can be destabilizing components of an N-degron^20^, we investigated sequence patterns with multiple bulky residues paired with a canonically stable P1 residue. We observed that sequences with bulky residues exclusively in P2/P3 had a significant drop in average PSI relative to the global population (ES = 0.25) (Figure 4D). In a deeper analysis of sequences with bulky P2 & P3 residues, we observed that the P1 residue with the lowest average PSI was Cys, followed by Gln and His (Figure S4B). Second, we observed that low PSI motifs with noncanonically destabilizing P1 often maintained a neutral to negative charge (Figure 4E). Interestingly, these low PSI sequences follow a charge profile that is opposite to that of the canonical six destabilizing P1 amino acids, where positively charged motifs appear more frequently at lower PSI. Third, the largest decreases in PSI were observed for canonically stable P1 sequences with both bulky and negatively charged residues (Figure 4F). These sequence dependent trends were not observed in the ClpS− dataset (Figure S4C). In total, these findings reveal a low stability series of sequences with P1 Cys, His, and Gln and bulky and negatively charged P2-P5 motifs that appear to be dependent on ClpS.

We next turned our attention to P1 Cys to understand what could make this synthetically useful P1 residue a component of an N-degron. Interestingly, P1 Cys had a lower mean PSI (3.42) than other stable P1 residues (PSI 3.80), suggesting that there are unstable sequences (Figure S2A). Upon visualizing the sequence composition of the 20,000 lowest PSI P1 Cys motifs, we observed an enrichment of bulky and negatively charged residues, whereas positively charged and flexible residues as well as P2 Pro were depleted in these motifs (Figure S4D). In a parallel analysis using ClpS− dataset, there was no enrichment of negatively charged sequences, suggesting that this was unique to ClpS expressing strains (Figure S4E). These enrichment trends aligned with residue-by-position PSI shifts for sequences with P1 Cys (Figure 2B, P1 Cys). We generated clonal isolates of five P1 Cys sequences with candidate destabilizing motifs. Two of the five candidates presented sfGFP:mCherry ratios on par with the FLFVQEL degron control, while the remaining three offered intermediate values (Figure S4F). When we performed a Western blot, at least one candidate sequence (CFYEAISDFI) was fully degraded, producing a non-visible band similar to the strongest identified degrons (Figure 4G).

We investigated other commonly functionalized P1 residues in Gly and Ser. Both residues were stable at P1, though an analysis of P2-P5 residues for the 20,000 lowest PSI sequences for P1 Gly and P1 Ser revealed an enrichment in Gln, Glu, Lys, and Trp at P2-P5 (Figure S4G & Figure S4H).

### Similar residue sidechains can cumulatively amplify or dampen bacterial N-degrons

We next studied the combinatorial effect of multiple amino acids, as stability can be a function of aggregate properties such as sidechain hydrophobicity or charge, or could otherwise be dictated by local sequence context similarly to other reported degrons^63^. To select which properties to target, we grouped amino acids based on their R-group characteristics and evaluated their enrichment in low stability sequences within the WT dataset. Unstable sequences showed an enrichment for aromatic, hydrophobic, and positively charged residues, and a deficiency in flexible, negatively charged, and small amino acids at various P1-P5 positions, but most distinctly at P1 and P2 (Figure 5A). Recognizing the differential enrichment in charged residues as well as literature precedent that penultimate charged residues can impact ClpSAP recognition and turnover^14,23,60^, we investigated whether there could be connection between the net charge of the N-terminal region and protein stability. We observed that lower PSI sequences are more common in neutral to positively charged motifs, whereas negatively charged motifs, even with P1 FLYWRK, had a significant increase in average PSI (Figure 5B). To further investigate, we tested clonal isolates of charged sequences carrying P1 Phe. We found FDEDE (−4) to be highly stabilizing and FDEAA (−2) to be lightly stabilized relative to a control degron, and that positively charged FRKAA (+2) was stabilized (Figure 5C). This finding is supported by the presence of negatively charged residues D36 and E41 flanking the N-degron binding pocket in ClpS crystal structure, which would repulse negatively charged P2-P5 substrates to make binding unfavorable (Figure S5A). Additionally, in the WT dataset both bulky (FLWY) and positively charged (RK) P1 residues showed increases in average PSI when paired with negatively charged P2-P5 motifs. P1 RK sequences demonstrated a higher increase in mean PSI between neutral and negatively charged P2-P5 datasets (Figure S5B & Figure S5C). This charge-based phenomenon was absent in the ClpS^−^ dataset (Figure S5D).

**Figure 5.**
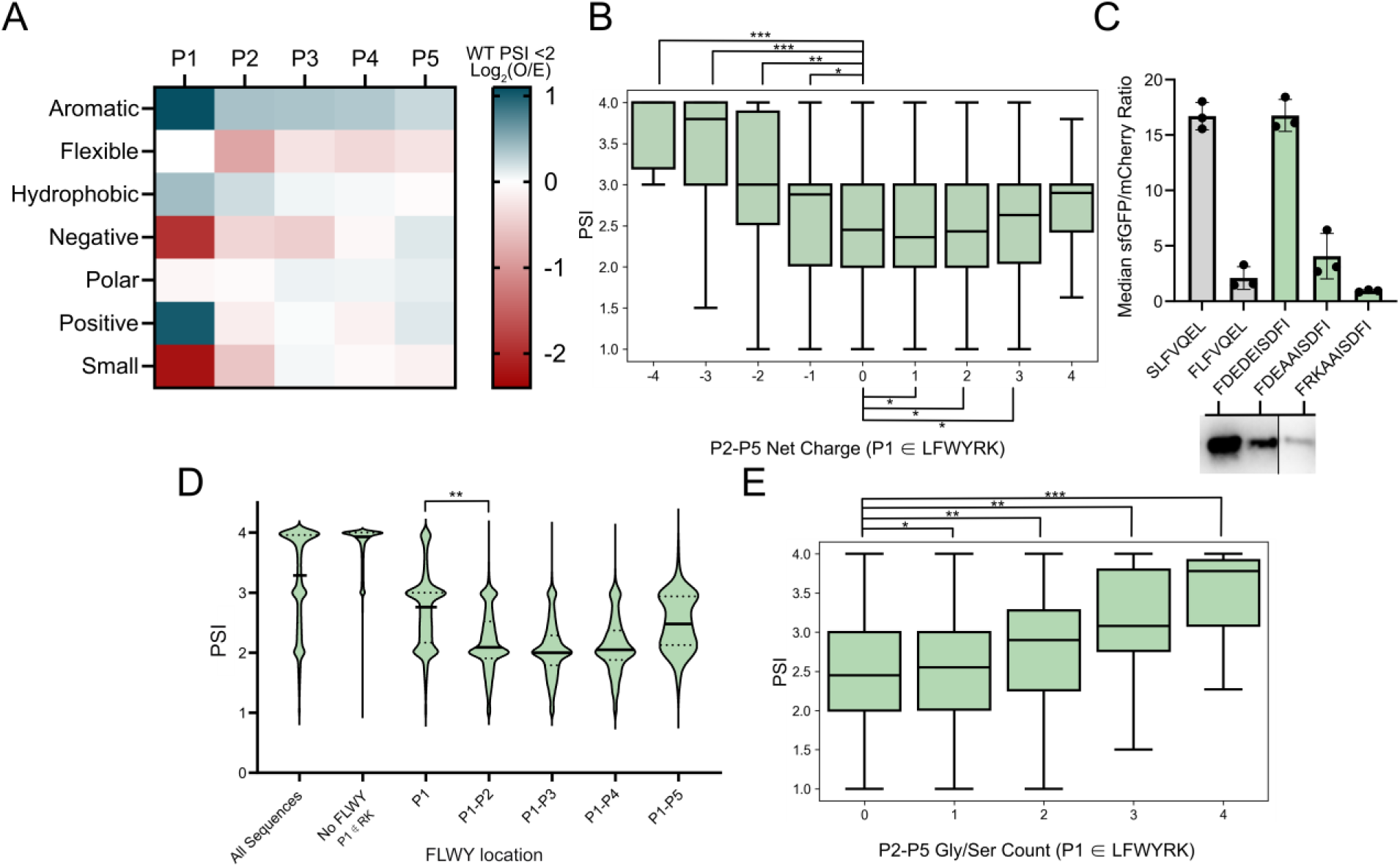
P1-P5 sidechain properties can cumulatively impact stability. **A**. Low stability motif heatmap depicting log_2_ enrichment (observed/entire dataset) of grouped amino acids. Negative = DE, Positive = RK, Hydrophobic = AVILMFWY, Polar = RNDQEKHSTYC, Aromatic = FWYH, Small = GASC, Flexible = GSDN. **B.** Boxplot showing that sequences with P1 FLWYRK and negatively charged P2-P5 show increased bulk stability. (Net charge, n, ES relative to 0 net P2-P5 charge): (−4, 35, 0.62), (−3, 1,341, 0.61), (−2, 22,647, 0.39), (−1,155,936,0.15), (0,399,998, N/A), (+1, 171, 933, 0.04), (+2, 30,185, 0.01), (+3, 2,422, 0.07), (+4, 75, 0.24). **C.** Negative charge can stabilize a canonically destabilizing P1 residue. (Top) Flow cytometry data for P1-P5 sequences with varying P2-P5 charge (FDEDE: −4, FDEAA: −2, FRKAA: +2). (Bottom) Western blots of corresponding sequences. Non-consecutive lanes from the same blot are separated by a solid line. **D.** Boxplot depicting that clustered FLWY at and near the N-terminus leads to decreased stability. P1(n=206,557) compared to P1+P2 (n=54,022) shows significance (p<0.05, ES = 0.45). **E.** Boxplot showing increased PSI means and distributions for sequences with P1 FLWYRK and increasing Gly/Ser residue counts in P2-P5. (Gly/Ser count, n, ES relative to 0 P2-P5 Gly/Ser): (0, 477,006, N/A), (1, 257,126, 0.07), (2, 46,829, 0.25), (3, 3,505, 0.50), (4,96, 72). All p-values in comparison to 0 P2-P5 Gly/Ser show p <0.05. p-values evaluated using a Mann-Whitney U test, effect size magnitude (ES) ranges from 0 to 1, as evaluated using Cliff’s delta. * = p <0.05, ES 0-0.25; ** = p <0.05, ES 0.25-0.50, *** = p <0.05, ES 0.50-0.75, **** = p <0.05, ES 0.75-1.0

We next analyzed the impact of clustered FLWY residues to investigate whether multiple bulky residues can additively impact stability. For sequences carrying multiple N-terminal FLWY residues, the average PSI was 0.56 units lower (p < 1E-99, ES = 0.45) for two such residues at P1 and P2 compared to a singular P1 FLWY. Diminishing PSI shifts were observed for motifs carrying >2 N-terminal FLWY residues (Figure 5D). On average, sequences carrying a single positively charged residue at P1 exhibited a lower average PSI than sequences with a bulky P1, and lower magnitude PSI shifts when multiple charged residues were chained (Figure S5E). However, these motifs may have multiple bulky residues appended onto P1 Arg or P1 Lys via LFTR polyaddition^17,18^. We screened several clonal isolates with multiple bulky and positively charged residues at P1-P5 and found them all to be destabilizing (Figure S5F).

We additionally analyzed the additive impact of multiple smaller flexible amino acids, focusing on Gly and Ser, which frequently compose flexible N-terminal linkers. In line with the depletion of small amino acids in destabilizing motifs, we observed PSI increase with increasing Gly/Ser count downstream of a P1 FLWYRK residue (PSI increases for Gly/Ser counts 0-1: 0.09, 0-2: 0.32, 0-3: 0.64, 0-4: 0.99, all p<1E-33) (Figure 5E). As such, the presence of these flexible motifs may aid in the stability of N-terminal motifs with canonically unstable residues.

### Training and validation of N-FIVE, an N-terminal stability predictive model

Our next objective was to develop a stability-predictive machine learning model to interpret the full sequence space. To generate a base matrix, we one-hot encoded each motif by amino acid and position. During training, we observed improved statistical performance upon applying a minimum read count threshold of 20 for any given amino acid motif. We applied this filter to trim our WT dataset to approximately 798,000 sequences prior to final training. We supplied these encodings and the corresponding PSI from 80% of the WT dataset to generate the training input. We then trained an extreme gradient boosted model using Python’s XGBoost library. After hyperparameter tuning, our model obtained an RMSE of 0.34 and an r-squared of 0.82 when validated using the remaining 20% of data. Satisfied with the performance of our model, we dubbed our model N-FIVE and set out to test its predictive capabilities (Figure 6A-6B). We first simulated the PSI of 100,000 random N-terminal sequences and observed the characteristic decrease in mean PSI for sequences carrying destabilizing N-termini (Figure 6C). Upon simulating the full 3.2M sequence space using N-FIVE, we compared its performance to our actual dataset, finding comparable P1 prediction and higher P2-P5 average PSI values (Figures S6A & S6B). We generated SHAP plots to understand the underlying architecture, observing trends on par with known N-degron pathway rules (Figure S6C).

**Figure 6.**
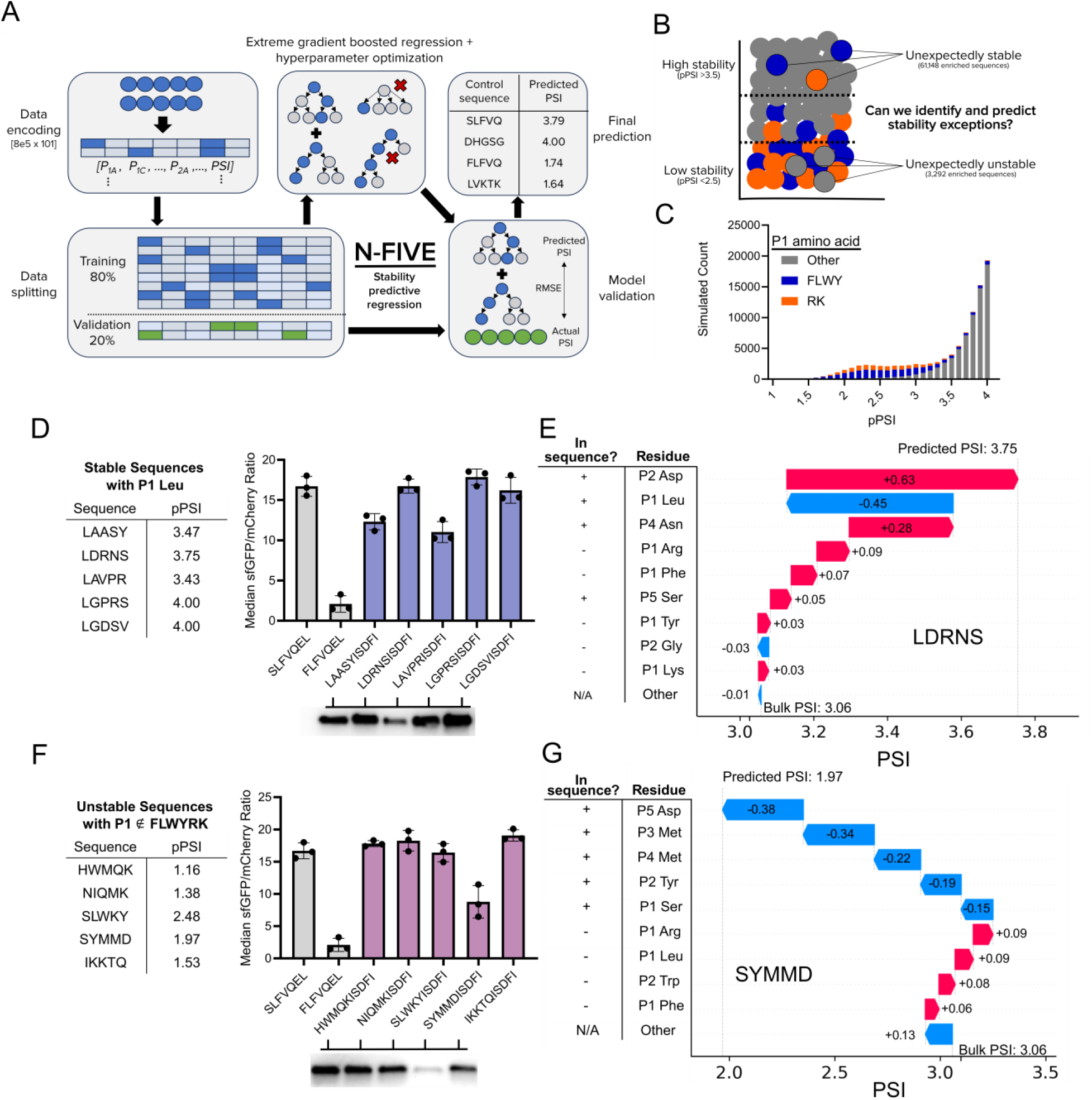
N-FIVE, a machine learning model for stability prediction. **A.** Architecture of the N-FIVE machine learning model. The first 5 amino acids of control sequences (Figure 1D) were predicted using N-FIVE and found to match the expected stability profile. **B.** High stability sequences (predicted PSI >3.5) are under-enriched in P1 LFWYRK and low stability sequences (predicted PSI <2.5) are over-enriched in P1 LFYWRK. **C.** PSI prediction of 100,000 randomly selected 5-amino acid sequences using N-FIVE. Sequences with a canonically destabilizing P1 are highlighted (bulky residues in blue, positively charged residues in orange). **D.** (Left) Predicted PSI values for five unexpectedly stable (pPSI >3.5) candidate sequences selected with a P1 Leu constraint. (Top right) Candidate sequence flow cytometry data, (right left) Western blots of the degron-sfGFP product. **E.** (right) SHAP values for individual matrix contributions to the predicted PSI value of 3.75 for the candidate sequence LDRNS. **F.** Predicted PSI values for five unexpectedly unstable (pPSI < 2.5) candidate sequences selected with a P1 ∉ FLWYRK constraint. (Top right) Candidate sequence flow cytometry data, (bottom right) Western blots of the degron-sfGFP product. **G.** (right) SHAP values for individual matrix contributions to the predicted PSI value of 1.97 for the candidate sequence SYMMD.

To explore the capabilities of N-FIVE, we studied N-terminal motifs with unexpected stability profiles. We sought to identify five stable sequences with a canonically destabilizing P1 Leu. Excitingly, all five of the sequences (sequence LXXXXISDFI) suggested by N-FIVE displayed stable profiles (Figure 6D). We additionally attempted the challenging task of identifying sequences that are unexpectedly unstable with P1 residues that are not in FLWYRK. From our five-sequence screen, one motif, SYMMDISDFI, generated a moderately destabilizing profile (Figure 6F). SHAP values were calculated for candidate sequences LAASYISDFI (Figure 6E) and SYMMDISDFI (Figure 6G), revealing the cumulative effect of P2-P5 residues in “overriding” the P1 residue.

## Discussion

This work presents the most comprehensive mapping of the *Escherichia coli* N-degron pathway to date. Having collected and analyzed over 2.2M screened sequences that cover the critical P1-P5 space within wild-type and adaptor deficient BL21, we both reinforce and provide insight into the *in vivo* substrate preferences of pathway with unprecedented granularity. Our findings corroborate the consensus understanding within *E. coli* that six amino acids present at P1 (Leu, Phe, Tyr, Trp, Arg, Lys) lead to lowered protein stability due to their recognition and degradation through the ClpSAP proteolytic cascade. We have additionally shown that stability determinants beyond P1 appear in our aggregation of screened sequences. Specifically, our analyses show that neutral to positively charged N-terminal motifs with P1 FLWYRK and local bulky and Gln residues are highly probable substrates for ClpS recognition. Frequently, P2-P5 bulky amino acids lower protein stability. However, it is unclear how they may be causing instability as there is conflicting evidence about whether bulky residue downstream of P1 can be bound directly in the hydrophobic pocket of ClpS^20^. It is possible that this instability could be caused by strengthened interactions in the substrate channel or surface of the protein. Furthermore, we show cases where the expected level of protein stability based on P1 can be fundamentally altered, such as when multiple small, stabilizing residues flank a canonically destabilizing P1 residue. The impact of penultimate residues is evident for destabilizing P2 Gln and P2-P5 bulky residues, the universally stabilizing P2 Pro and Gly, sequence-specific stabilization from internal Asp, Glu, and Ala, and the scattered P2-P5 preferences for P1 FLWY, amongst other examples.

These findings bear relevance to various applied settings, including bacterial N-degradation, N-terminal binders, and the design of synthetic N-termini for applications in small molecule/protein ligation in live cells. For example, bioconjugation efforts that rely on a P1 Cys should avoid bulky and negatively charged residues near the N-terminus. Gly-Ser motifs are likely to provide a stabilizing effect that would offset any degradation-promoting elements. For N-terminal identification, the low tolerance for negatively charged motifs may be a challenge. The mutation and screening of variants that have lowered negative electrostatic potential near the binding pocket may address this limitation. When tuning protein expression using degrons, the characteristic structure of an N-degron mentioned in the previous paragraph is worth considering when designing constructs with a target stability in mind.

Amongst the most destabilizing motifs were those that could be generated by LFTR. Multiple previous studies have identified that polyaddition of Leu and Phe, rather than addition of a singular Leu or Phe, is a possibility^17,18^. The highly destabilizing nature of multiple Leu/Phe-Arg residues in tandem coincide with previous observations that the turnover of LR-substrates is greater than just Leu-substrates.^14^ Furthermore, others have suggested that the ClpS binding site has been optimized to accommodate LFTR-modified substrates^67^. On balance, the generation of Leu/Phe-rich motifs in front of Nt-Arg and Lys may contribute to the generation of highly destabilizing degrons, supporting the rapid turnover of LFTR-substrates despite the requirement of an additional modification.

Interestingly, the P2-P5 profile for P1 Cys degrons aligns with that of the eukaryotic Ac/N-degron pathway, where the N-acetyltransferase NatA acetylates N-terminal cysteine with a preference for polar and acidic residues downstream of P1 Cys^68^. Within bacteria, N-acetylation has been observed by GNAT-family enzymes RimI-RimJ, including for Nt-Cys^69^, but it has not has not yet been linked to protein degradation. As such, the exact mechanism for the instability of these sequences is unknown. Our data does suggest a ClpS-mediated phenomenon, yet there may be an uncharacterized protein that modifies these motifs.

For the synthetic design of N-terminal motifs, the stability-predictive N-FIVE model can be consulted to evaluate or suggest motifs even when there are sequence constraints. The predictive power of N-FIVE has been evidenced by its ability to solve a complex sequence-stability problem; in this instance successfully suggesting unexpectedly stable sequences. The identification of unexpectedly unstable sequences remains a challenge, in part due to the small number of candidate sequences to form the dataset. Further development of more complex ML models may improve stability prediction capabilities. As such, to aid in such analyses, we have made our full datasets and trimmed, storage friendly sequence-stability databases available for public use. Finally, our application of SHAP values supports a thought process for evaluating the strength of an N-degron; one where a given sequence is measured as accumulation of stabilizing or destabilizing factors that build to a net predicted effect.

## Acknowledgements

The authors thank the following people: Dr. Arit Ghosh of the University of Delaware Flow Cytometry Core for training and advice on cell sorting and analysis. Bruce Kingham and Mark Shaw of the University of Delaware Sequencing & Genotyping Center for consultations and performing NGS sequencing. Ishika Govil and Zhao Wei Wang for discussions and code for the analysis of NGS data. Prof. Karl Schmitz for discussions regarding ClpSAP interactions and manuscript review. Prof. Kevin Solomon for discussions regarding bioinformatic analysis. We further thank members of the Kunjapur Lab for their helpful discussions and review of the manuscript.

## Author Contributions

S.S. designed and performed most experiments, wrote the bioinformatics and machine learning code, analyzed data, and wrote the manuscript. N.C. assisted in optimizing DNA library generation. A.T. assisted in developing NGS data analysis code. K.K. assisted with cloning and assay development. A.M.K. oversaw the project, designed experiments, and assisted in writing and editing the manuscript.

## AI Statement

Bioinformatics analysis code was developed and debugged with the assistance of ChatGPT.

## RESOURCE AVAILABILITY

### Lead contact

Further information and requests for resources and reagents should be directed to and will be fulfilled by the Lead Contact, Aditya Kunjapur.

### Materials availability

This study did not develop new unique reagents or chemicals.

Plasmids and strains are available from the Lead Contact upon request.

### Data and code availability

NGS analysis code, trimmed sequence databases, and the N-FIVE model have been made available at GitHub (https://github.com/KunjapurLab/N-terminal-cluster-stability). The raw FASTQ.gz datasets will be made publicly available by the time of publication. Uncropped blots and raw data can be found in the supplemental files.

## EXPERIMENTAL MODEL AND SUBJECT DETAILS

*E. coli* DH10β and *E. coli* E. cloni SUPREME were used for plasmid construction and *E. coli* BL21 (DE3), *E. coli* BL21*clpS-*, and *E. coli* BL21 *aat-* sequence screening.

## Methods

### EXPERIMENTAL MODEL AND STUDY PARTICIPANT DETAILS

#### Microbial culture

*Escherichia coli* strains, including BL21, ClpS-BL21, and LFTR-BL21, were routinely grown in culture tubes containing LB media at 37 °C while shaking at 250 RPM, unless mentioned otherwise.

### METHOD DETAILS

*Escherichia coli* strains and plasmids used are listed in Table S1 and Table S2, respectively. *clpS* and *aat* inactivations were performed as described in the Genetic knockout of N-recognins section. Genes were purchased as G-Blocks or gene fragments from Integrated DNA Technologies (IDT) or Twist Bioscience and were optimized for E. coli K12 using the IDT Codon Optimization Tool. A version of scUbp1 without the N-terminal region was used for improved expression in *E. coli*^70^. Essential genetic sequences and primers can be found in Table S3 and Table S4, respectively. Cloning amplicons were generated using KOD XTREME Hot Start polymerase and corresponding buffering reagents. Amplicons were verified and purified by running through a 1% agarose gel for 200 V for 20 minutes followed by gel excision and extraction. Samples were assembled in homemade Gibson assembly aliquots run at 50 °C for 30 minutes. Assemblies were transformed into *E. coli* cloning strains DH5α or DH10β using either electroporation or chemical heat shock. Following a one-hour outgrowth after transformation, cells were plated on permissive antibiotic plates and were incubated at 37 °C overnight. The following afternoon, multiple colonies were picked and incubated overnight in separate culture tubes containing LB media (10 g/L tryptone, 5 g/L sodium chloride, 5 g/L yeast extract) and the permissive antibiotic. The next day, cells were stocked 1:1 with 30% glycerol at negative 80 °C for future usage. The remaining cells were miniprepped for sequencing. Plasmid sequences were verified through a mixture of Sanger sequencing through Eurofins, Genewiz, and Azenta and full plasmid sequencing through Plasmidsaurus. Plasmids were transformed into BL21 and BL21 derivatives using either electroporation or chemical heat shock. Two plasmid systems were transformed in series.

#### Materials and chemicals

The following compounds were purchased from MilliporeSigma (Burlington, MA, USA): phosphate-buffered saline (PBS), kanamycin sulfate, glycerol, 25 nm membranes (VSWP02500) and KOD XTREME Hot Start polymerase (Millipore 71975-3). D-glucose was purchased from TCI America (Portland, OR, USA). Agarose, Laemmli SDS sample reducing buffer, and ethanol were purchased from Alfa Aesar (Ward Hill, MA, USA). Anhydrotetracycline (aTc) was purchased from Cayman Chemical (Ann Arbor, MI, USA Methanol, sodium chloride, LB Broth powder (Lennox), LB Agar powder (Lennox), Amersham ECL Prime chemiluminescent detection reagent, and Thermo Scientific™ Spectra™ Multicolor Broad Range Protein Ladder were purchased from Fisher Chemical (Hampton, NH, USA). Taq DNA ligase was purchased from GoldBio (St. Louis, MO, USA). Phusion DNA polymerase and T5 exonuclease were purchased from New England BioLabs (NEB) (Ipswich, MA, USA). SybrSafe DNA gel stain was purchased from Invitrogen (Waltham, MA, USA). E. cloni 10G Supreme electrocompetent cells were purchased from Biosearch Technologies. Miniprep kits were purchased from Zymo. 4-20% precast protein gels were purchased from Bio-Rad. Flow cyometry filter caps (Chemglass Life Sciences CLS4380009), arabinose, DpnI enzyme (FERFD1704) and Immobilon-E Western blot membranes were obtained from Fisher Scientific. HRP-conjugated anti-6*His antibody (Proteintech HRP-66005) was obtained from Proteintech (Rosemont, IL, USA).

#### Genetic knockout of N-recognins

Multiplex automatable genome engineering (MAGE) was used to inactivate the endogenous *aat* (LFTR) and *clpS* genes^59^. MAGE oligonucleotides were designed using MODEST^71^ to insert three in-frame stop codons into the gene of interest. Freshly made electrocompetent cells were resuspended in 5 μM oligonucleotide and subsequently electroporated to enable cell permeation of the oligonucleotide. Cells were outgrown in LB media for an hour prior to plating on permissive plates. To verify the desired genetic knockouts, allele-specific colony PCR was performed using KAPA 2G Fast HotStart polymerase (Roche KK5801). Additional Sanger sequencing was performed to verify asPCR hits.

#### Fluorescence analysis of degrons

Overnights made from biological triplicate colonies of BL21 and recognin-deficient BL21 derivatives carrying both the dual reporter and Ubp1 expressing plasmids were inoculated at a 1:100 ratio of grown culture : fresh LB media containing 0.2% arabinose, 100 ng/mL aTc, 20 μg/mL chloramphenicol, and 15 μg/mL kanamycin. Cells were grown at 37 °C, 250 RPM for 18 hours prior to analysis. Cells were then filtered using 35 μm filter caps into cytometry tubes containing fresh PBS at a roughly 1:250 ratio of cell culture: PBS. Cells were analyzed on a NovoCyte Flow Cytometer (Agilent Technologies). Prior to analysis, scatter gates were used to isolate singlet bacterial cells of the appropriate morphology. sfGFP fluorescence was analyzed using a 488 nm laser and a 530/30 nm bandpass filter and mCherry fluorescence was analyzed using a 488 nm laser and a 660/20 nm bandpass filter. For all analyses, at least 100,000 events that passed through the singlet gate were collected.

#### Western blotting of hexahistidine-tagged reporter proteins

1 mL of overnighted culture containing expressed dual reporter and Ubp1 protease were lysed using glass beads for 15 minutes. The sample was transferred to a fresh microcentrifuge tube and centrifuged at 12,000 RPM for 10 minutes. Subsequently, the supernatant was collected. Sample concentration was evaluated using a Bradford assay on a SpectraMax i3x with a BSA calibration curve. Sample concentration was normalized to 0.1 mg/mL in water with 1x SDS PAGE loading dye and denatured at 95 °C for 10 minutes. To separate proteins by size, 10 uL of sample is loaded into a 4–20% Mini-PROTEAN TGX gel and run at 180 V for 35 minutes. Subsequently, the proteins were transferred onto an Immobilon-E membrane through an overnight wet transfer (25V for 15 hours on ice). The following day, the membrane was blocked for at least one hour with 5% milk in TBST buffer. Subsequently, an HRP-conjugated anti-His6x antibody was added to the blocking solution at a 1:10,000 dilution, and was allowed to incubate at room temperature for 1 hour. Following three washes in TBST to remove the solution, ECL Prime chemiluminescent reagent was added to the membrane. Following a 10 second incubation, chemiluminescent images were captured at varying intensities using an Azure c280.

#### DNA library preparation

DNA library plasmids were designed to be assembled from two separate PCR amplicons. Primers containing 5 consecutive NNK codons at P1-P5 of the degron template were ordered from IDT, using a 25A/25C/25G/25T (N) or 50G/50T (K) ratio of hand-mixed nucleotides to maximize diversity in the mutagenized region. PCR amplicons were generated using KOD XTREME polymerase using roughly 200 ng of plasmid template per reaction with an annealing temperature of 59 °C and extension time of 2 minutes for inserts and 3 minutes for the backbone. To maximize sequence diversity and avoid overamplification, 5 separate PCR amplicons were generated at a low cycle count (20 cycles) and high volume (50 μL per reaction). Amplicons were then digested using a DpnI enzymatic digestion to remove the template. The amplicons were then isolated on a 1% agarose gel and subsequently purified using gel extraction. Amplicons were pooled and assembled using an isothermal Gibson assembly method with a total of 300 ng of total DNA at a 1:1 molar ratio of insert: backbone. Assemblies were then dialyzed on a 25 nm membrane placed over deionized water for 60 minutes. Five separate assemblies were pooled and then electroporated into E. cloni 10G Supreme electrocompetent cells. Immediately following electroporation, cells were mixed with 500 μL of SOC media and were placed in a culture tube in a 250 RPM shaking incubator set to 37 °C. Following a 30-minute outgrowth, cells were plated at serial dilutions to verify transformation efficiency. For 20^5^ member libraries, a transformation efficiency of 10^8^ colonies/transformation was the minimum threshold established for further experimentation. Following a second 30-minute outgrowth, 5 mL of SOC media was added to the tube and cultures were allowed to grow overnight.

The following morning, library stocks were made by mixing 500 μL of undiluted culture at a 1:1 ratio with 30% glycerol, which were then stored at −80 °C indefinitely. The remaining 4-4.5 mL of culture was miniprepped using a Zymo ZR Plasmid Miniprep-Classic kit. BL21 and recognin-deficient BL21 derivatives carrying a Ubp1-expressing plasmid were made electrocompetent using standard molecular biology procedures, concentrating 10 mL cultures to 200 μL competent cell aliquots. Miniprepped libraries were electroporated into freshly made competent cells. Outgrowth, efficiency measurements, overnight growth, and freezer stocking were performed similarly to original library transformations. A transformation efficiency of 10^8^ was obtained for all conditions. Library diversity was confirmed prior to sorting using 2.5 Mbp next-generation sequencing through Azenta’s AmpliconEZ platform.

#### Cell culturing and fluorescence activated cell sorting

Overnights of BL21 and recognin-deficient BL21 derivatives carrying both library plasmids and Ubp1-expressing plasmids were inoculated at a 1:100 ratio in fresh LB media containing 0.2% arabinose, 100 ng/mL aTc, 20 μg/mL chloramphenicol, and 15 μg/mL kanamycin. Cells were grown at 37 °C for 18 hours prior to analysis. Cells were then filtered using 35 μm filter caps into cytometry tubes containing fresh PBS at a roughly 1:250 ratio of cell culture: PBS. Cells were sorted on a BD FACS Aria Fusion using a 70 μm or 85 μm nozzle and a neutral density 1.0 filter. Prior to analysis, scatter gates were used to isolate singlet bacterial cells of the appropriate morphology. sfGFP fluorescence was analyzed using a 488 nm laser and a 530/30 nm bandpass filter and mCherry fluorescence was analyzed using a 561 nm laser and a 610/20 nm bandpass filter. Bins were set based on controls, where SLFVQEL sorted into B4 and template RCGGAIISDFI sorted into B2 (Supplemental Figure 1). A small gap was placed between bins to maximize sort efficiency, and all bins maintained a sort efficiency ≥ 95%. Sorting conditions and sample concentrations were optimized to obtain event rates on the order of 1,000 – 5,000 events per second. 10 million gated cells were sorted into bins covering the sfGFP range, excluding the bottom of the distribution that was observed to have a higher concentration of debris and GFP-inactivating mutants. The following cell counts were collected in the format [B4; B3; B2; B1]. WT: [5,998,196; 1,940,219; 1,130,665; 955,183]. LFTR-: [6,953,470; 1,161,141; 998,851; 913,809]. ClpS-: [7,048,173; 1,651,034; 483,574; 853,250]. Cells were recovered overnight in a mixture of 1:1 recovered cells: LB media containing 1% glucose. The subsequent day, cells were stocked and miniprepped using methods described in the DNA library preparation section.

#### Next generation sequencing preparation and analysis

250 base pair (AVITI) and 405 base pair (Amplicon EZ) DNA amplicons for next-generation sequencing were designed to contain a 5’ and 3’ sequencing adapter, a 3 base pair barcode for demultiplexing, and a centralized mutagenized region to ensure coverage by both paired-end reads. The full sequence can be found in our corresponding methods paper^61^. Amplicons were generated using KAPA 2G polymerase, minimizing cycle counts (20 was default) to decrease over-enrichment of select sequences. The amplicons were then isolated on a 1% agarose gel and subsequently purified using gel extraction. Samples were then submitted to Azenta’s Amplicon EZ service (∼100,000-300,000 2×250 paired end reads) or to the UD Sequencing & Genotyping Center (>100 million 2×150 paired end reads on an Element AVITI). The WT, LFTR-, and ClpS− datasets had 146,704,252, 180,209,552, and 188,333,530 reads respectively for B1-B4 and the presorted library. All sequencing datasets had a Q30 >95%.

NGS data was first converted from a FASTQ.gz format to FASTA for ease of analysis. When necessary, reads were demultiplexed using the three base pair barcodes that corresponded to unique sorted bins. Reads were compared to the consensus sequence for the submitted amplicon. Any reads that varied from the consensus sequence (excluding the barcode and mutagenized region) or those that contained stop codons (TAG, TAA, TGA) in the mutagenized region were then excluded from subsequent analysis. The mutagenized region was identified in each read based on a 15 base pair consensus sequence located directly after the mutagenized region, and the mutagenized nucleotide sequence was translated into an amino acid sequence. From data collected for each strain, read counts were measured for each five amino acid motif across the four bins. From this data, the protein stability index (PSI) was calculated using the formula 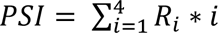, where *R_i_* is the fraction of reads present in bin *i*^48^. Each unique sequence, its read counts per bin, and PSI were stored in a database for subsequent analysis.

Heatmaps, boxplots, and violin plots were visualized using the Seaborn and matplotlib Python packages^72,73^ as well as GraphPad Prism. Weblogos were generated using Logomaker^74^. All code was developed using Python. Code, sequence databases, and N-FIVE are accessible on GitHub: https://github.com/KunjapurLab/N-terminal-cluster-stability. Full NGS data will be made available prior to publication.

#### Development of N-FIVE machine learning model

During early rounds of model training, we observed that applying a 20 read minimum filter to included sequences minimized RMSE and improved R^2^ for the model. As such, data from the wild-type BL21 sequence-PSI database were one-hot encoded into an 798,837 x 101 matrix. The first 100 columns represented a binary encoding of each of the 20 amino acids across 5 positions, while the final column corresponded to the PSI value. The dataset was randomly split, with 80% of the data being used for model training and 20% being used for model validation and evaluation. An extreme gradient boosted nonlinear regression model was then trained using the Python XGBoost^75^ package for 20 rounds. This type of model outperformed random forest regressors and ridge regression models statistically. Hyperparameter optimization was performed using Optuna^76^ over 30 trials, aiming to minimize RMSE. The following parameters were set: *n_estimators*: 986, *max_depth*: 11, *learning_rate*: 0.085, *subsample*: 0.904, *colsample_bytree*: 0.810, providing an RMSE of 0.34 for predicted PSI. The final model was then saved as a .pkl file for future analysis. To evaluate the contribution of each matrix component, SHAP (Shapley Additive exPlanations) values were calculated^77^ for the whole model as well as for individual sequences.

### QUANTIFICATION AND STATISTICAL ANALYSIS

To quantify differences in PSI distributions between defined groups, we applied two non-parametric statistical methods: the Mann-Whitney U test and Cliff’s Delta for effect size calculations. The Mann-Whitney U test was selected due to the skewed nature of PSI distribution as well as its robustness to unequal sample sizes. We used a two-sided Mann–Whitney U test to assess whether the PSI distributions between two groups differed significantly. Cliff’s Delta was computed to quantify the magnitude and direction of the distributional shift between compared groups. Unlike common effect size measurements such as Cohen’s d, Cliff’s Delta makes no assumptions about the underlying distribution of PSI values and remains valid for skewed, bounded, or ordinal data. Here, Cliff’s Delta represents the probability that a randomly selected value from one group is greater than a randomly selected value from the other, minus the reverse probability to yield an effect size bounded between −1 and +1.

For general enrichment analysis, the counts of any given residue or motif were first measured. To calculate enrichment differences between two populations, the average occurrence of an amino acid or motif in the appropriate population was calculated and subsequently subtracted to obtain a difference. To obtain a ratio, groupings were normalized to the same sample size and subsequently divided for ratiometric comparison. Log_2_ analysis was performed at times to simplify data visualization.

For clonal isolate testing, fluorescence measurements were taken for at least 100,000 gated cells using flow cytometry using cultures grown from three separate biological triplicates. Median ratiometric fluorescence ratios +/− standard deviation were reported. Median values were selected over mean values due to large mean shifts observed in select cases due to outliers, to which median values were less sensitive.

## Supplemental Content

Document S1. Figures S1–S6, Tables S1 & S2

Document S2. Uncropped Western blots presented in Figures 3-6

---

**Supplemental File 1.** Supplemental Figures 1-6 and Supplemental Tables 1-2

**Supplemental Figure 1.**
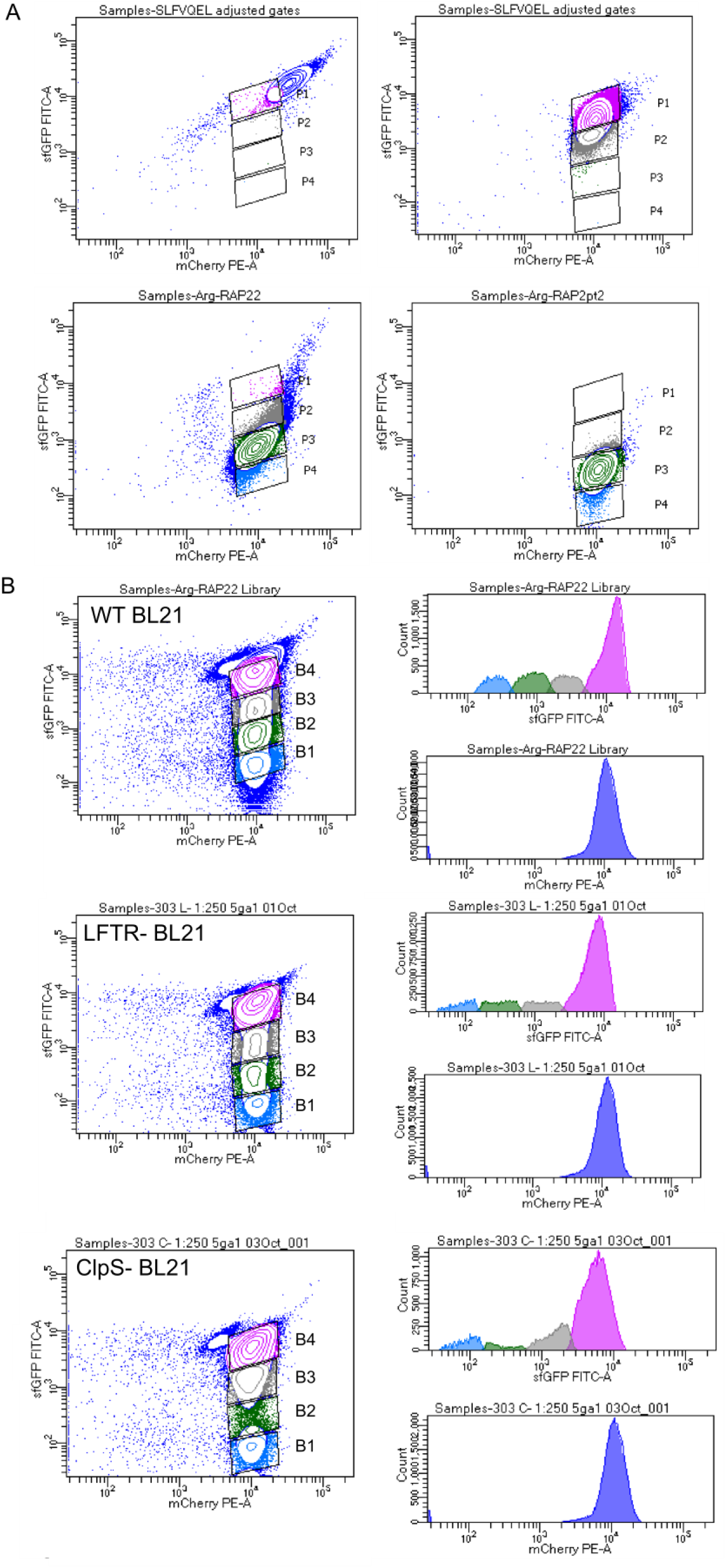
A.) Differentiation of stable/unstable sort controls. SLFVQEL (stable, top) and RCGGAIISDFI (unstable, arginylated library template, bottom) sequences hardcoded into the dual fluorescent reporter are differentiated across two separate examples on a BD FACS Aria Fusion. Left samples were collected prior to the WT 10M event sort. B.) 10M event sort information. sfGFP (y-axis) vs. mCherry (x-axis) distributions for the sorted WT (top), LFTR^−^ (middle), and ClpS^−^ (bottom) libraries with corresponding bins (left). Histograms for sfGFP & mCherry fluorescence (right).

**Supplemental Figure 2.**
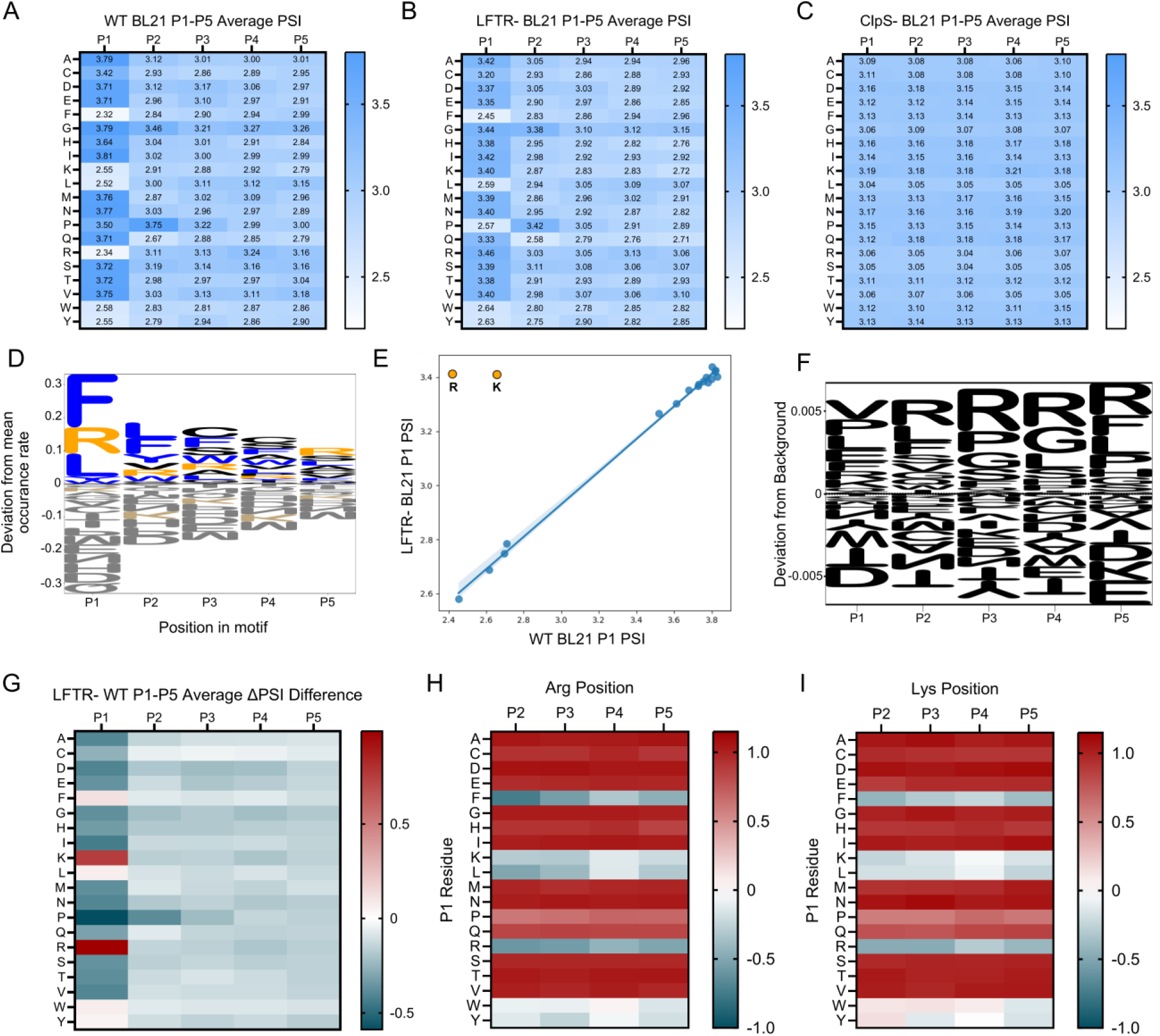
A.) P1-P5 WT BL21 Heatmap A mean PSI heatmap generated from 2.29 million sequences for the first five positions of a protein in wild-type BL21. B.) P1-P5 LFTR-BL21 Heatmap A mean PSI heatmap generated from 2.34 million sequences for the first five positions of a protein in BL21 deficient in aat (LFTR). C.) P1-P5 ClpS-BL21 Heatmap A mean PSI heatmap generated from 2.19 million sequences for the first five positions of a protein in BL21 deficient in clpS. D.) Bulky and positively charged amino acids are enriched in low stability sequences. Weblogo depicting the difference in enrichment for each amino acid in each of the five positions in low PSI (PSI <2) samples compared to the entire dataset. E.) P1 amino acid correlation between WT BL21 and LFTR- BL21 sorts Correlation between PSI scores for N-terminal residues between WT BL21 and LFTR- BL21 (r = 1.00) in the 10 million event sort. Arg/Lys/Pro were excluded from regression analysis. Arg and Lys separately highlighted in orange. Ubiquitin upstream of P1 Pro is not fully cleaved, complicating analysis. F.) Pro, Arg, Leu, and small amino acids are weakly enriched in low PSI sequences in the ClpS− dataset. Enrichment values of amino acids in sequences with a PSI <2 in the ClpS− dataset were subtracted from enrichment values from the whole dataset. Changes in representation cumulatively were <1% in either direction. G. P1 Arg and P1 Lys stability shifts are observed as a primary difference between LFTR- and WT datasets. Heatmap depicts the change in PSI for each AA-position combination between LFTR- and WT datasets. H. Internal Arg heatmaps per P1 residue. Heatmaps depict the average PSI for Arg at each position for any given P1 residue in the WT dataset. I. Internal Lys heatmaps per P1 residue. Heatmaps depict the average PSI for Lys at each position for any given P1 residue in the WT dataset.

**Supplemental Figure 3.**
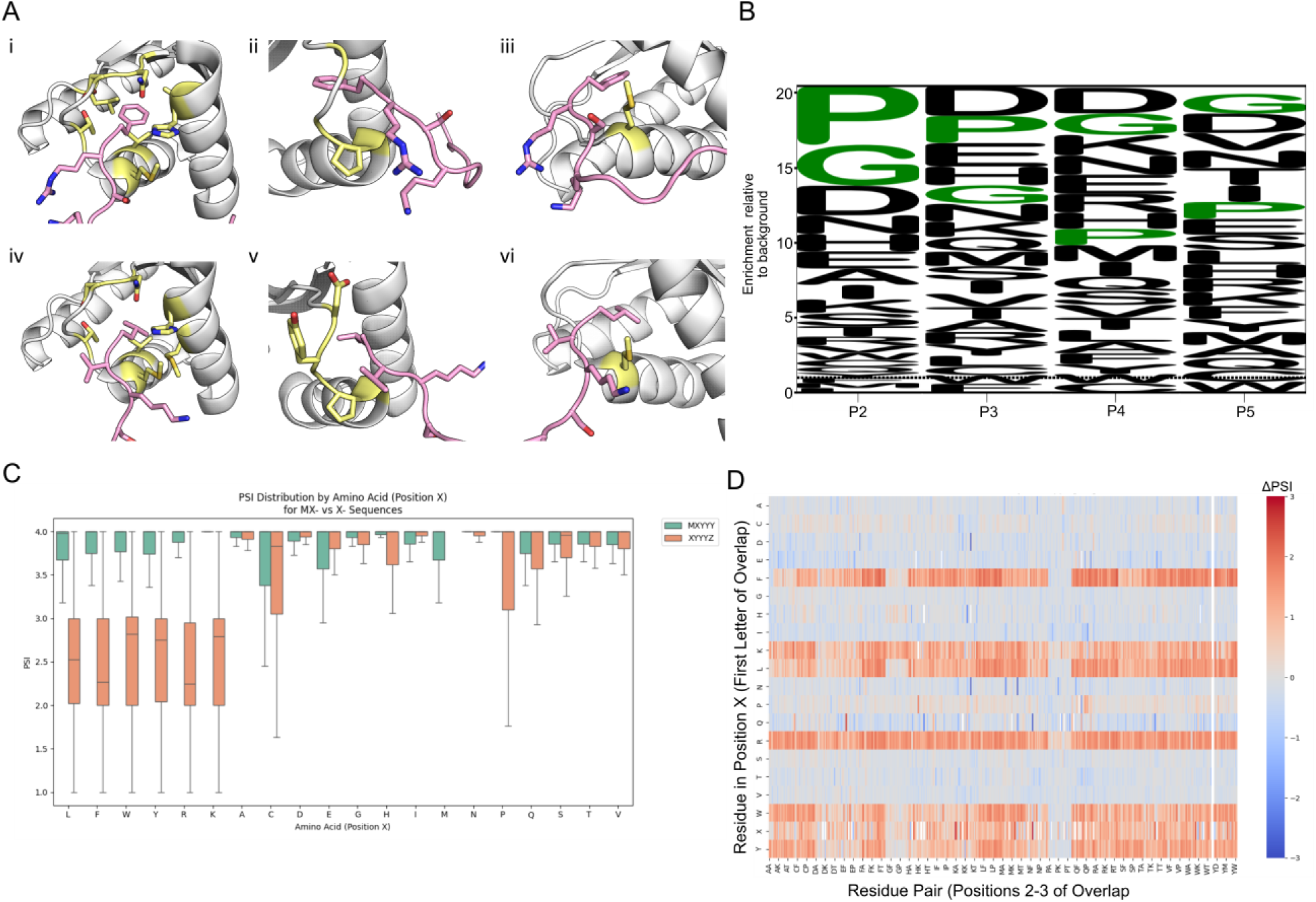
A.) Structures of ClpS bound to peptide substrates provide limited insight into the molecular basis of substrate specificity downstream of P1. i-iii: Structures of ClpS bound to Fpep with nearby ClpS residues highlighted in yellow for (i) P1, (ii) P2, and (iii) P3. (PDB ID: 2WA8) iv-vi: Structures of ClpS bound to Lpep with nearby ClpS residues highlighted in yellow for (iv) P1,(v) P2, and (vi) P3. (PDB ID: 2W9R). No interacting residues within 4 A were obtained for P4 and P5 of the peptides. B.) Glycine and proline are highly enriched in high stability P1 FLWYRK sequences. A P2-P5 WebLogo showing high stability P1 FLWYRK (PSI >3) sequence enrichment relative to the entire wild-type dataset for each of the five positions studied. Glycine and proline highlighted in green. C.) MX-Motifs do not show similar PSI distributions for P1 LFWYRK motifs. Mean PSI boxplots for residues in position X from the 10M event sort. MXYYY is in green and XYYYZ is in orange. D.) Change in PSI between MXYYY and XYYYZ datasets, with the X-position on the y-axis and the first two Y positions in the x-axis.

**Supplemental Figure 4.**
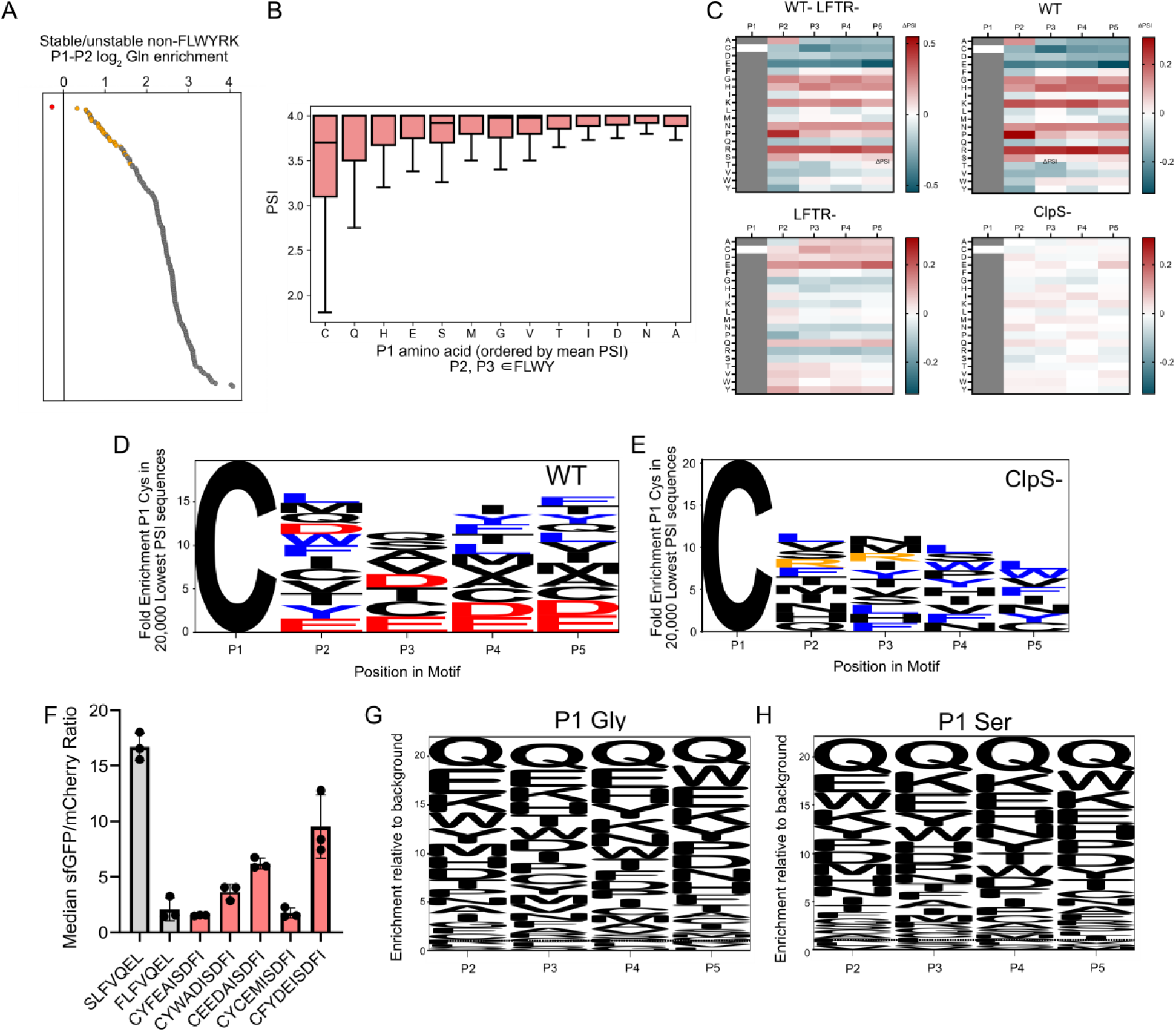
A.) P1/P2 Gln is noticeably enriched in low stability P1-P2 motifs that do not have P1 FLWYRK. Log2 enrichment of P1-P2 motifs in low stability (PSI <2) motifs relative to high stability (PSI >3) motifs where FLWYRK are excluded from P1. Motifs with Gln in P1 or P2 are in orange, QQ is in red. B.) P1 Cys is the most destabilizing non FLWYRK residue when P2 and P3 are bulky. Boxplot showing PSI distributions for non-P1 FLWYRKP motifs that have P2 and P3 residues in FLWY. C.) P1 Cys mean PSI heatmaps in various backgrounds reveal P2-P5 charge and bulky residue preferences. Sequences with P1 Cys were filtered in each dataset. Data represents differences from the P1 Cys global mean. WT- LFTR- represents the subtraction of difference from the mean values between the WT and LFTR- datasets. D.) Low stability P1 Cys sequences are rich in negatively charged and bulky amino acids at P2-P5. Weblogo depicting enriched amino acids for the 20,000 lowest PSI motifs with P1 Cys in the WT dataset. Bulky residues (FLWY) depicted in blue and negatively charged residues (DE) depicted in red. E.) Negatively charged residues are absent in low stability P1 Cys sequences. Weblogo depicting enriched amino acids for the 20,000 lowest PSI motifs with P1 Cys in the ClpS− dataset. Bulky residues (FLWY) depicted in blue and postively charged residues (RK) depicted in orange. F.) Full panel P1 Cys screen reveals putative N-degrons. Flow cytometry data collected from biological triplicates for the full panel of Nt-Cys screened sequences. G.) Enriched residues in low stability sequences for P1 Gly. Weblogo depicting the enrichment of amino acids relative to the whole population for the lowest 20,000 PSI sequences with P1 Gly. H.) Enriched residues in low stability sequences for P1 Ser. Weblogo depicting the enrichment of amino acids relative to the whole population for the lowest 20,000 PSI sequences with P1 Ser.

**Supplemental Figure 5.**
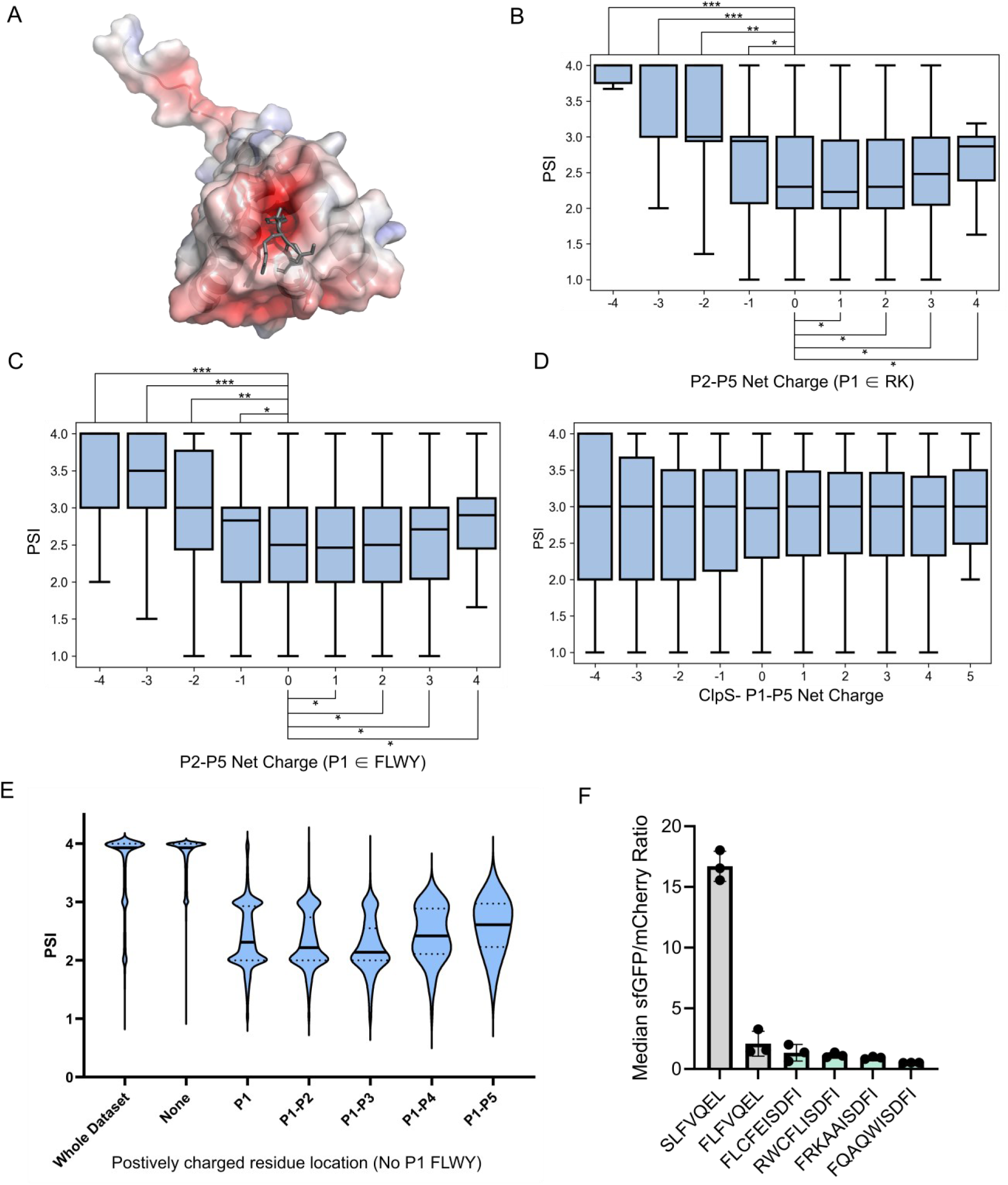
A.) The ClpS binding pocket has negative electrostatic potential. ClpS bound to a substrate (PDB 2WA8) visualized using the APBS Pymol plugin to create a surface map where regions of negative electrostatic potential are highlighted in red and regions of positive electrostatic potential are highlighted in blue. B.) Separation of charged and bulky P1 residue net charge heatmaps. C.) ClpS-sequences show charge parity. P1-P5 net charge heatmap for data collected in ClpS− Bl21. D.) Multiple positively charged residues at and near P1 lead to low PSI sequences. Violin plot depicting the PSI distribution in wild-type BL21 for sequences with one or more RK residues in series at and adjacent to the N-terminus. E.) Sequences with multiple FLWYRK(Q) residues show low stability. Flow cytometry data from the protein stability assay depicting sequences carrying multiple FLWYRK or P2 Gln residues exhibited low sfGFP:mCherry ratios relative to literature controls. All p-values in comparison to 0 P2-P5 Gly/Ser show p <0.05. p-values evaluated using a Mann-Whitney U test, effect size magnitude (ES) ranges from 0 to 1, as evaluated using Cliff’s delta. * = p <0.05, ES 0-0.25; ** = p <0.05, ES 0.25-0.50, *** = p <0.05, ES 0.50-0.75, **** = p <0.05, ES 0.75-1.0

**Supplemental Figure 6.**
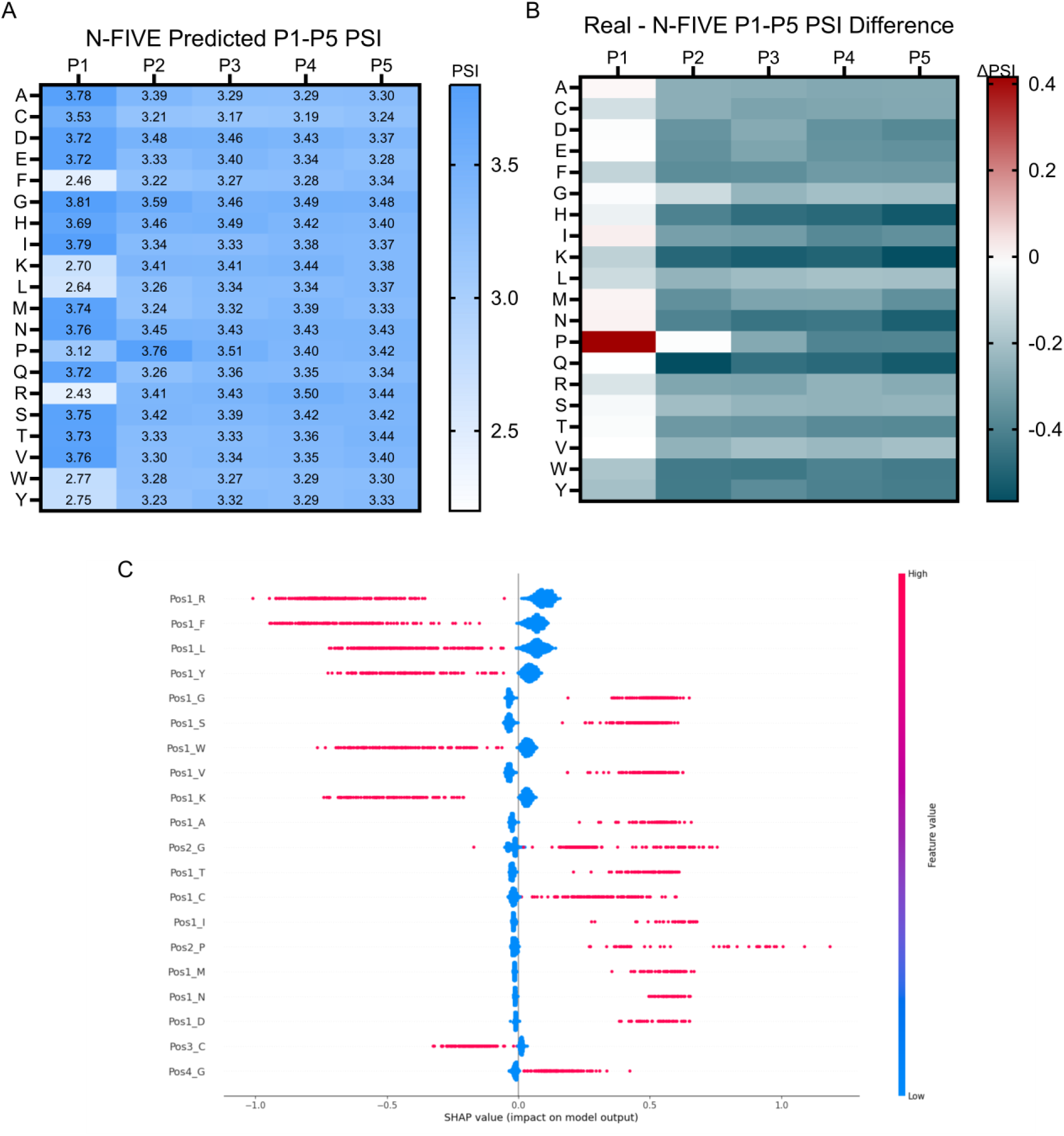
A.) N-FIVE PSI predictions for the mean PSI heatmap for all 3.2 million sequences in wild-type *Escherichia coli* BL21. B.) Change in PSI for each cell between the collected WT BL21 dataset and the N-FIVE predicted dataset. C.) Individual matrix components for N-FIVE predictions. SHAP values showing the contribution of the 20 most impactful matrix features onto the N-FIVE model.

**Supplemental Table 1.**
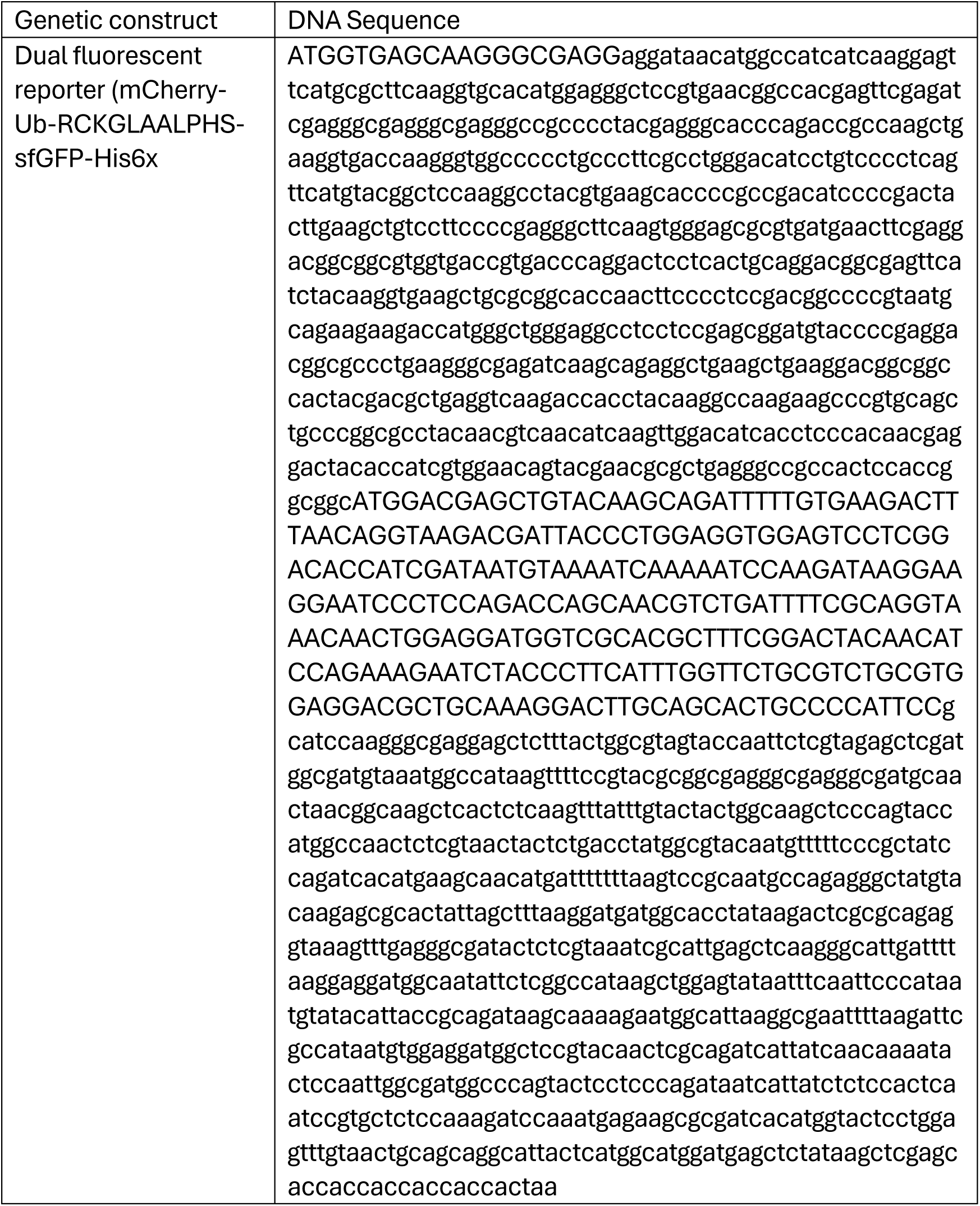
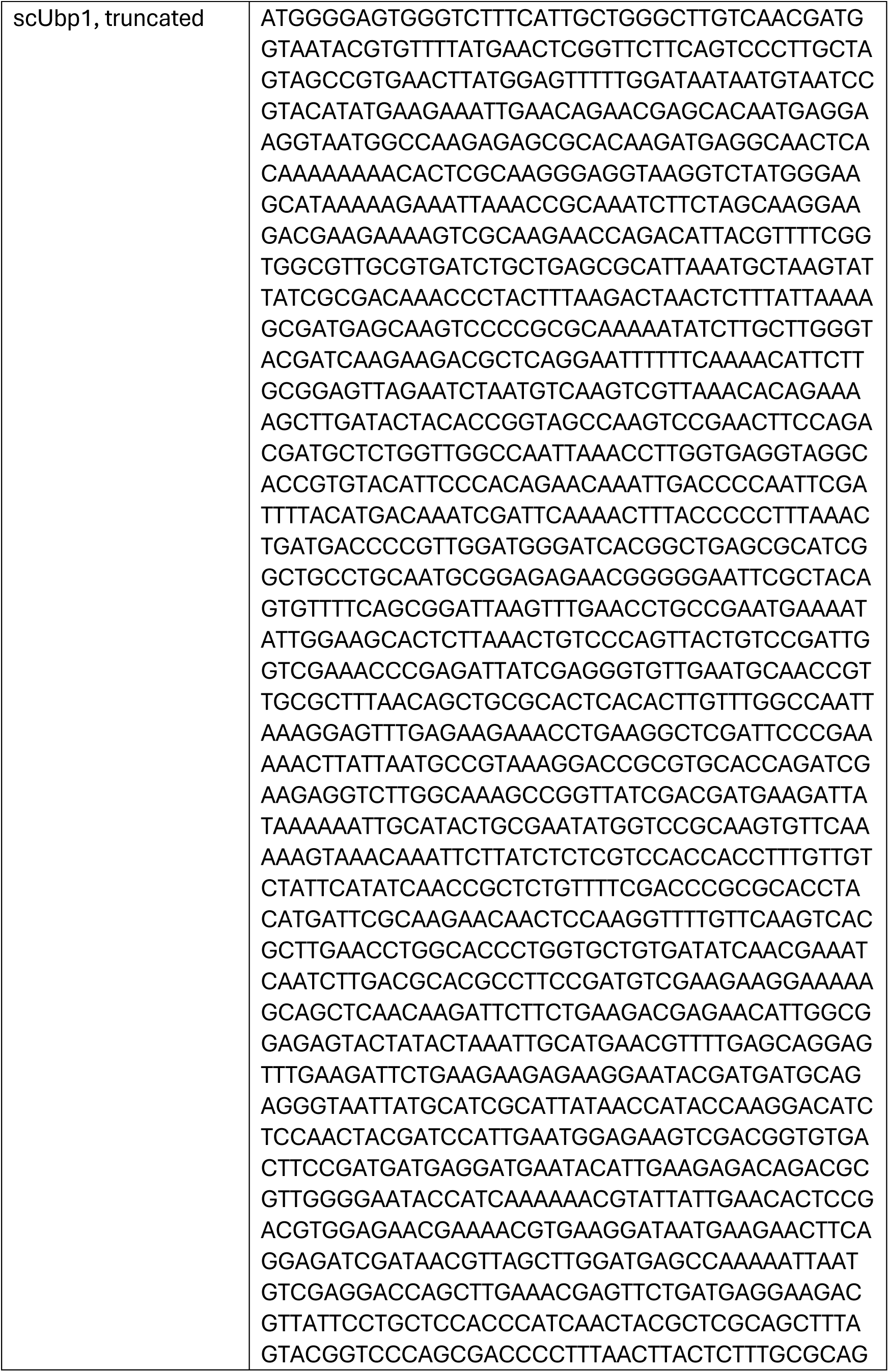
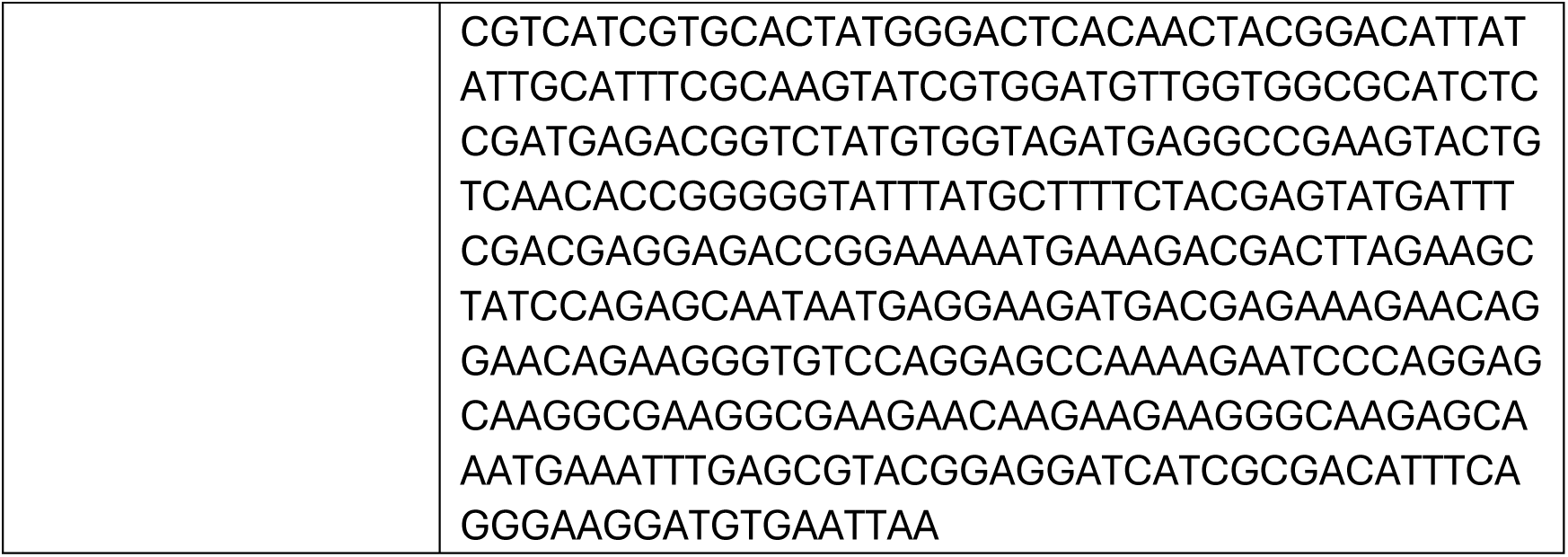
Important gene sequences.

**Supplemental Table 2.**
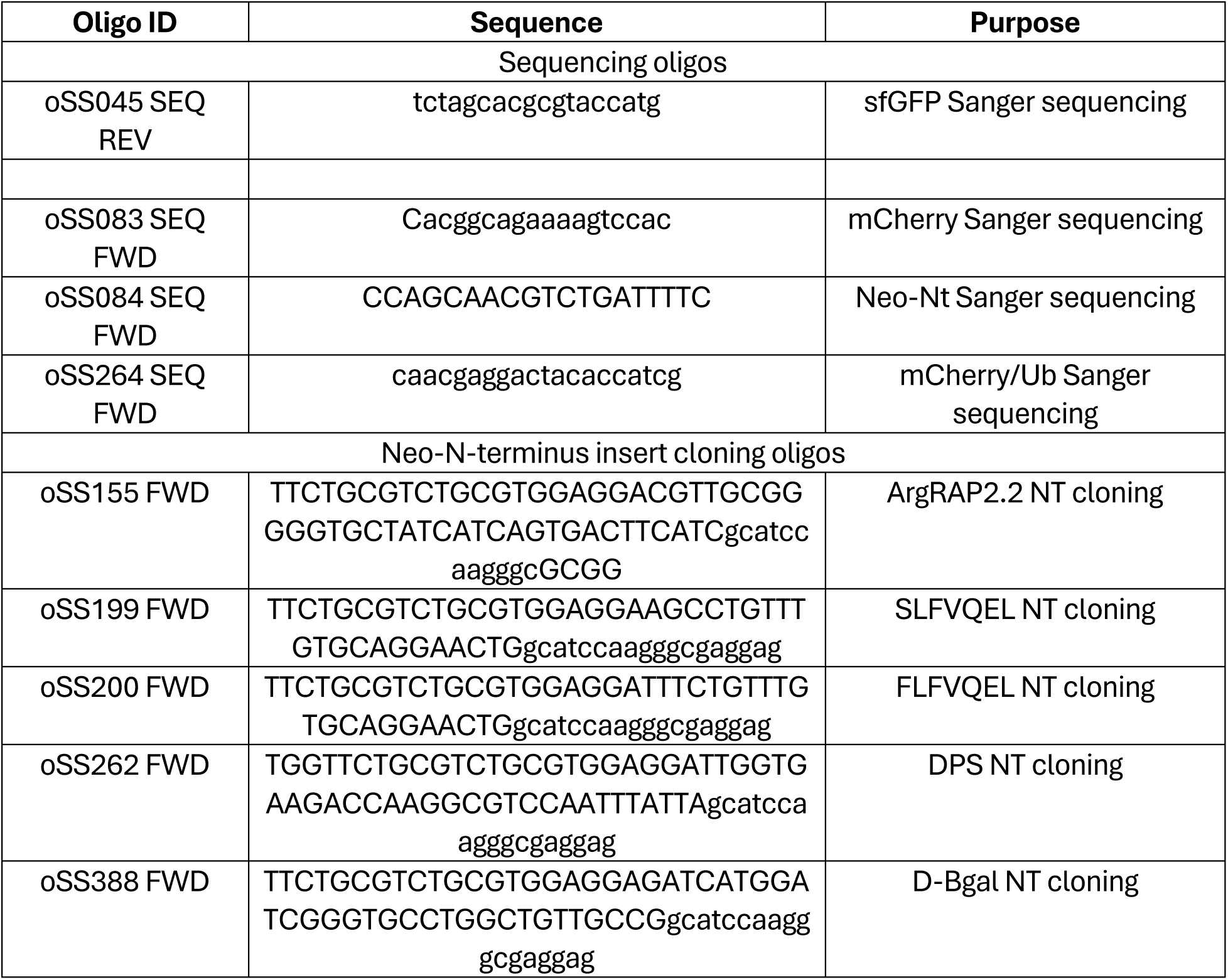
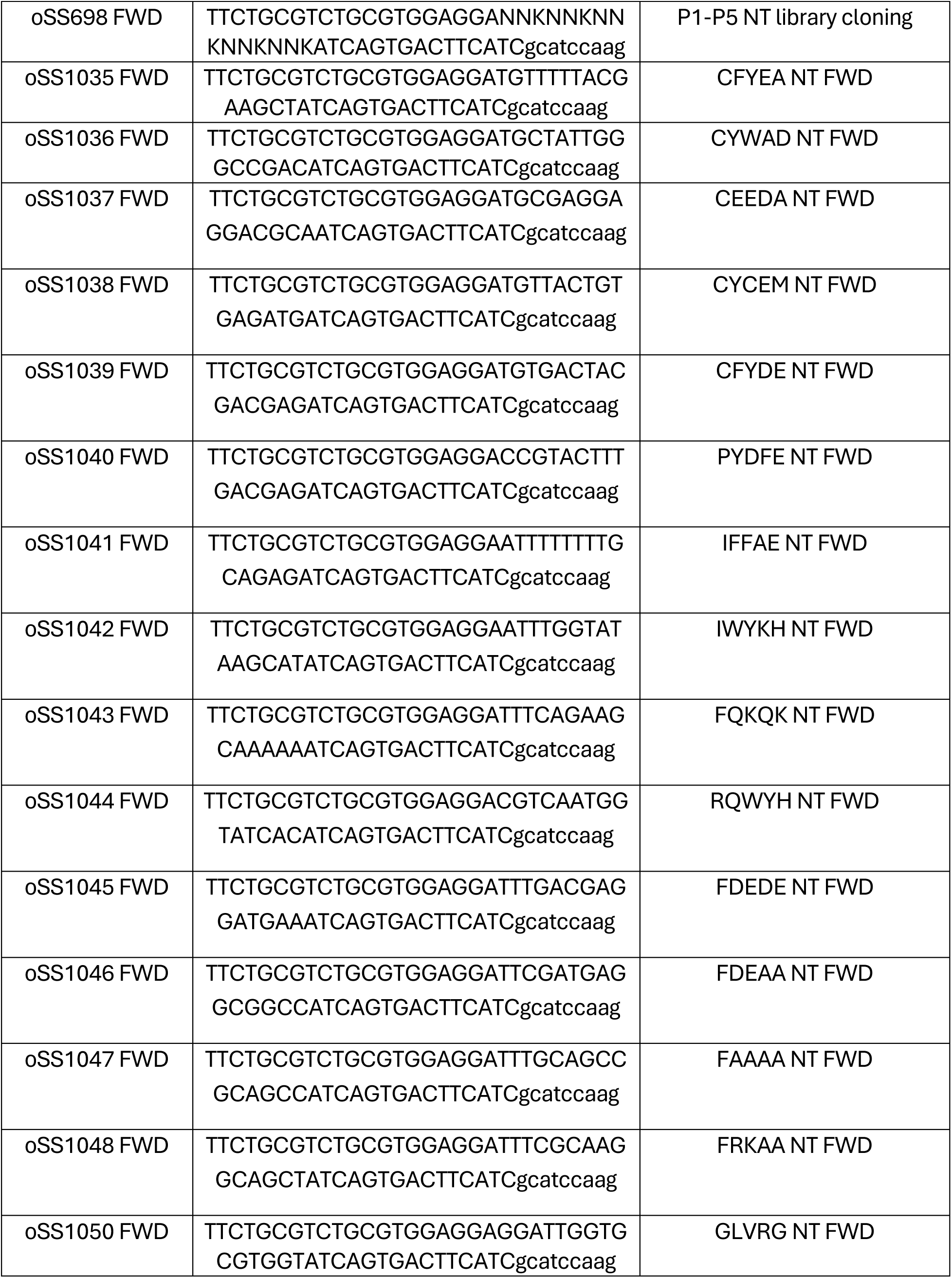
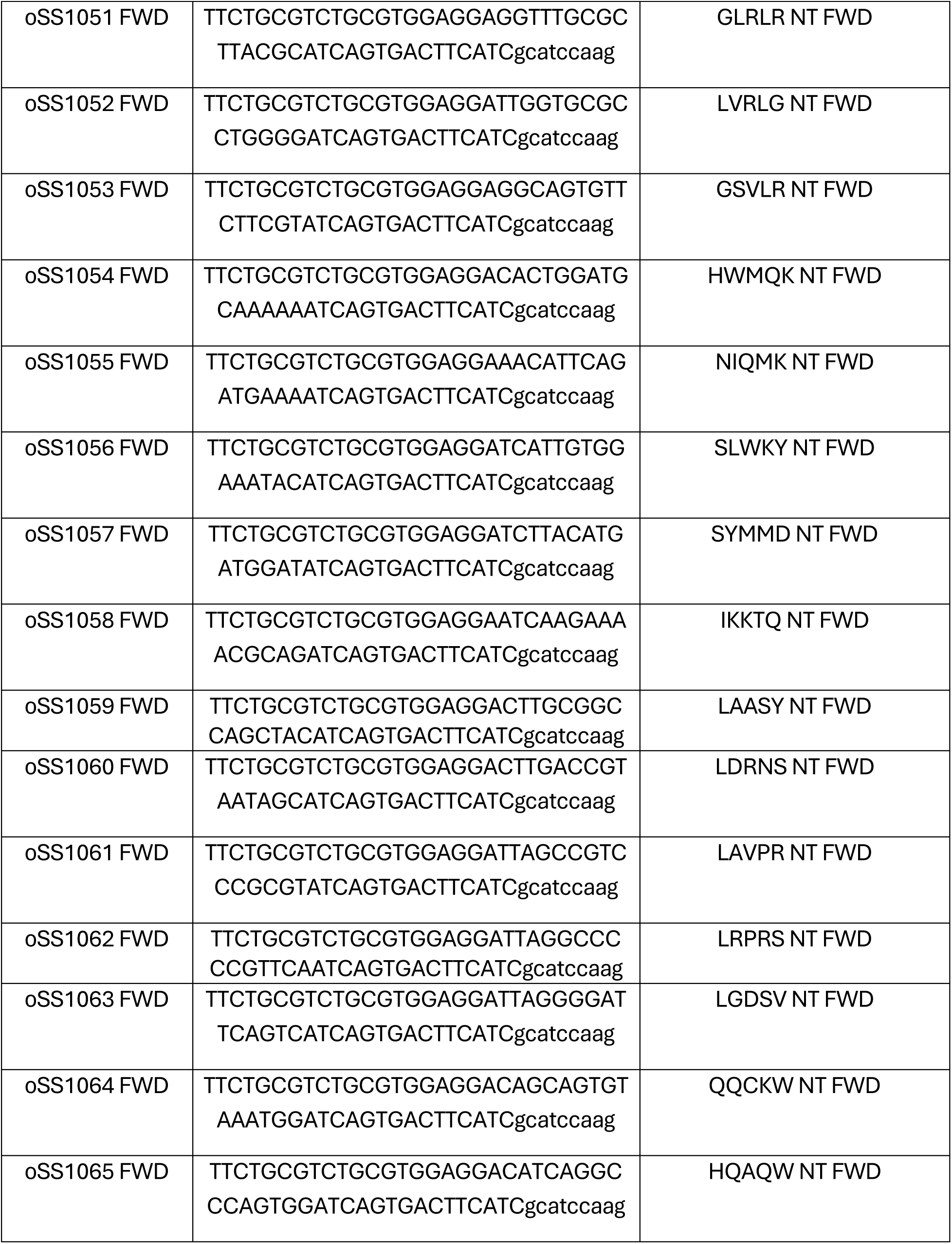
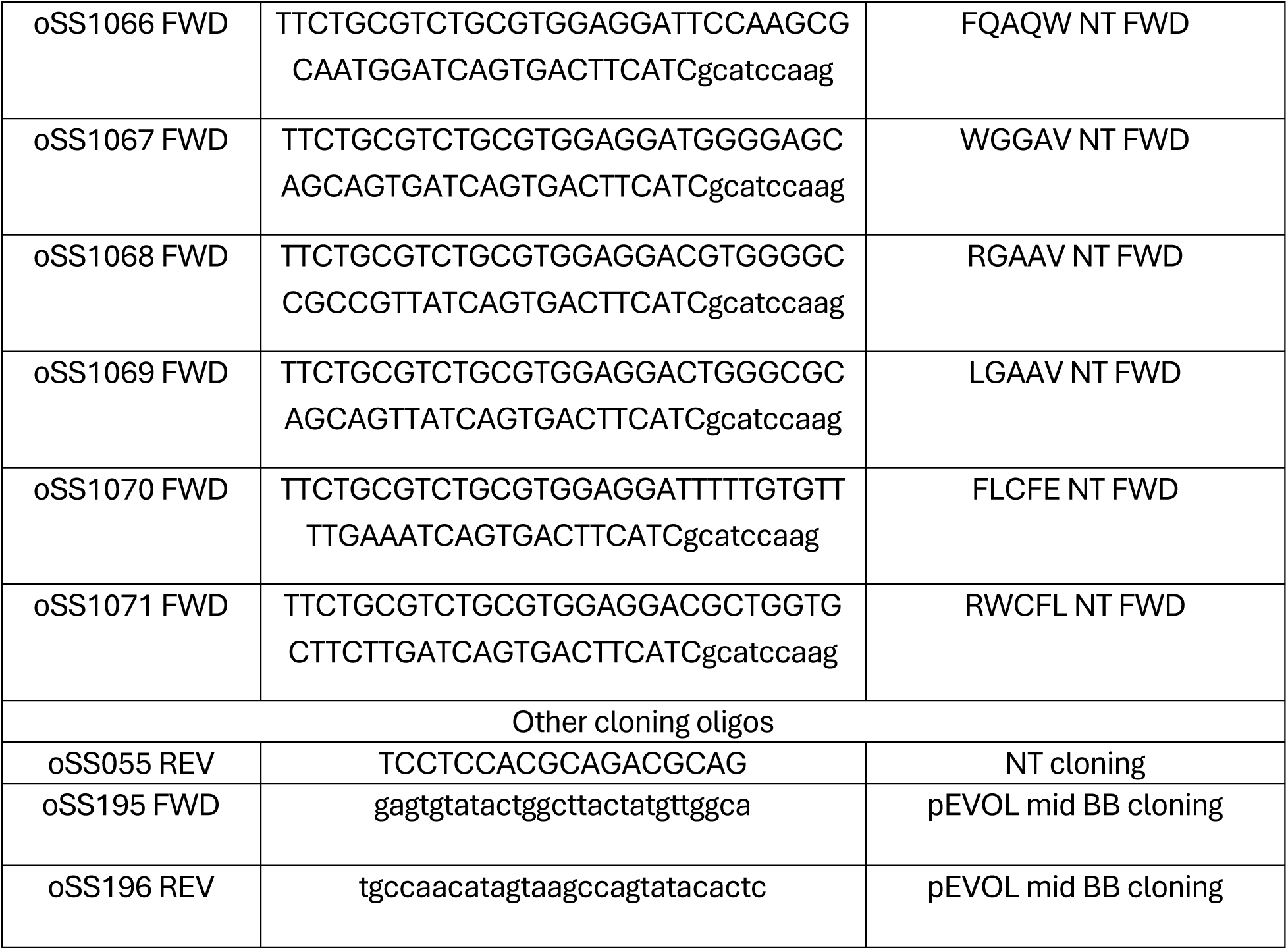
Important oligos. Oligos marked with SEQ were used for Sanger sequencing. Neo-N-terminus (NT) cloning featured a target NT primer + oSS196 as piece 1 and oSS195 + oSS055 as piece 2 used to generate amplicons from a pEVOL dual reporter plasmid.

**Supplemental File 2.**
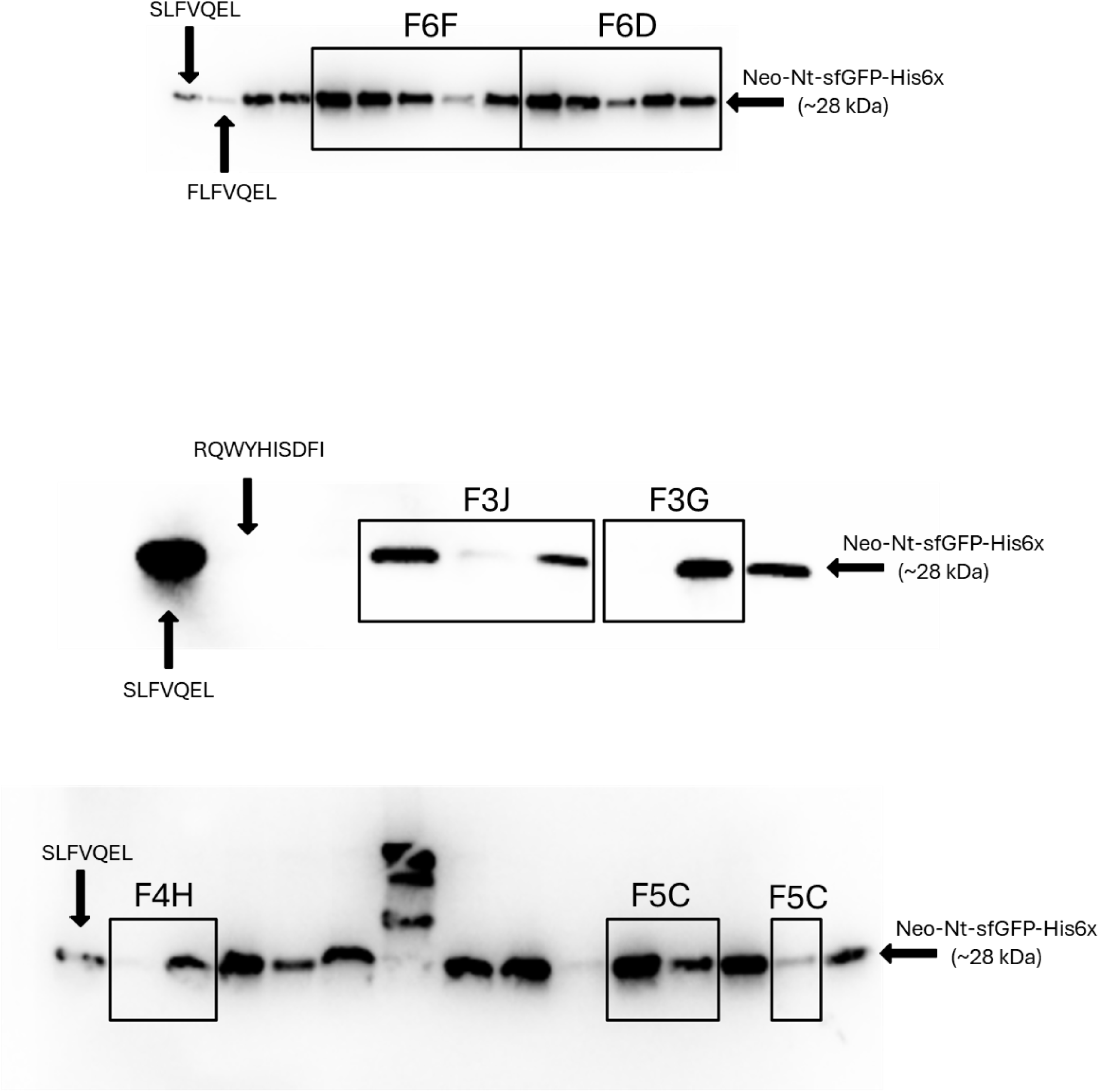
Uncropped Western blots. A consecutive lane was skipped in Figure 5C due to compounding factors that may promote stability for a neutrally charged motif, such as the presence of repetitive small amino acids at P2-P5 (FAAAISDFI).

